# Integrative single-nucleus multi-omics analysis prioritizes candidate *cis* and *trans* regulatory networks and their target genes in Alzheimer’s disease brains

**DOI:** 10.1101/2023.05.12.540037

**Authors:** Julia Gamache, Daniel Gingerich, E. Keats Shwab, Julio Barrera, Melanie E. Garrett, Cordelia Hume, Gregory E. Crawford, Allison E. Ashley-Koch, Ornit Chiba-Falek

## Abstract

**Background:** The genetic underpinnings of late-onset Alzheimer’s disease (LOAD) are yet to be fully elucidated. Although numerous LOAD-associated loci have been discovered, the causal variants and their target genes remain largely unknown. Since the brain is composed of heterogenous cell subtypes, it is imperative to study the brain on a cell subtype specific level to explore the biological processes underlying LOAD.

**Methods:** Here, we present the largest *parallel* single-nucleus (sn) multi-omics study to simultaneously profile gene expression (snRNA-seq) and chromatin accessibility (snATAC-seq) to date, using nuclei from 12 normal and 12 LOAD brains. We identified cell subtype clusters based on gene expression and chromatin accessibility profiles and characterized cell subtype-specific LOAD-associated differentially expressed genes (DEGs), differentially accessible peaks (DAPs) and *cis* co-accessibility networks (CCANs).

**Results:** Integrative analysis defined disease-relevant CCANs in multiple cell subtypes and discovered LOAD-associated cell subtype specific candidate *cis* regulatory elements (cCREs), their candidate target genes, and *trans*-interacting transcription factors (TFs), some of which were LOAD-DEG, for example, *ELK1* in excitatory neurons (Exc1) and *KLF13* and *JUN*, found in multiple cell subtypes. Finally, we focused on a subset of cell subtype-specific CCANs that overlap known LOAD-GWAS regions and catalogued putative functional SNPs changing the affinities of TF motifs within LOAD-cCREs linked to LOAD-DEGs including, *APOE* and *MYO1E* in a specific subtype of microglia and *BIN1* in a subpopulation of oligodendrocytes.

**Conclusions:** To our knowledge, this study represents the most comprehensive systematic interrogation to date of regulatory networks and the impact of genetic variants on gene dysregulation in LOAD at a cell subtype resolution. Our findings revealed crosstalk between epigenetic, genomic, and transcriptomic determinates of LOAD pathogenesis and define catalogues of candidate genes, cCREs, and variants involved in LOAD genetic etiology and the cell subtypes in which they act to exert their pathogenic effects. Overall, these results suggest that cell subtype-specific *cis-trans* interactions between regulatory elements and TFs, and the genes dysregulated by these networks contribute to the development of LOAD.

## INTRODUCTION

Large multi-center genome-wide association studies (GWAS) have identified associations between numerous genomic loci and late-onset Alzheimer’s disease (LOAD) ^1–7^. The most recent GWAS meta-analyses reported 75 risk loci for LOAD ^8, 9^. However, the precise disease-causing genes, the specific causal genetic variants, and the molecular mechanisms mediating their pathogenic effects have not yet been explained. Most LOAD-GWAS variants are in noncoding genomic regions^10^. Previous studies suggested that some noncoding LOAD SNPs are located in regulatory elements such as enhancers, affecting their functions, and thereby impacting gene expression ^11^, including expression of distal genes ^12^. Thus, to untangle the genetic and genomic architecture of LOAD and to translate LOAD genetic association discoveries to causal mechanisms of disease, LOAD GWAS variants need to be assigned to the correct target genes, rather than merely the nearest gene ^12, 13^. Moreover, it is imperative to map these variants and their linked genes to the specific brain cell-type in which they exert their pathogenic effect.

Many functional genomic studies provide evidence for the role of gene dysregulation in LOAD pathogenesis, including those examining specific disease-related genes ^14, 15^, pathways ^16^, expression quantitative trait loci (eQTLs) ^17–19^, differential transcriptome profiles^20^, and the DNA methylation ^21–24^ and histone mark ^25^ landscapes in human brain tissues. However, these studies were conducted in bulk brain tissue, and therefore cannot specify the cell type(s) in which gene expression or epigenetic changes occur. Mixed cell subtypes could also mask signals corresponding to a particular cell subtype, especially if the causal cell subtypes comprise a small fraction of the entire sample. Furthermore, bulk tissue studies are confounded by sample-to-sample variation in cell type composition, which could be exacerbated by the neuronal loss and proliferation of glial cells accompanying LOAD ^26^. Transcriptomic and epigenomic studies using sorting techniques to separate broad cell types from LOAD brain tissue ^27–32^ for example, neuronal vs. non-neuronal, have provided new important insights, but even within these categories, there are many different cell types and subtypes in the human brain ^33^. Single-cell experimental approaches can circumvent these limitations and inform LOAD-specific epigenomic and transcriptomic changes with unparalleled precision.

The past three years have seen a transition into single-cell multi-omics studies in LOAD functional genomic research. Single-cell transcriptomic studies have achieved previously unattainable resolution in identifying LOAD-associated cell type-specific changes in gene expression ^34, 35^. These studies demonstrate the importance of examining gene dysregulation at the cell type-specific level. However, they focused only on single-nucleus (sn)RNA-seq data and therefore the underlying regulatory mechanisms for these gene expression signatures remain to be identified. More recently, integrative multi-omics single-nucleus framework studies profiling chromatin accessibility and gene expression identified cell-type-specific, disease-associated candidate *cis*-regulatory elements (cCREs) and their candidate target genes, and demonstrated the utility and potential of this strategy in moving LOAD genetic research forward ^36, 37^.

The diagram in Figure 1a presents an overview of this study pipeline. We used the 10X Genomics platform to perform *parallel* snRNA-seq and snATAC-seq analyses simultaneously from the same pool of nuclei derived from post-mortem temporal cortex tissue of LOAD patients and neuropathologically normal controls. We used these datasets to identify LOAD-associated differentially expressed genes (DEGs), differentially accessible peaks (DAPs) and *cis* co-accessibility networks (CCANs) at the cell subtype level. The *parallel* experimental design allowed us to integrate these LOAD profiles to promote the mechanistic understanding of gene dysregulation in LOAD. We identified LOAD-associated cell subtype specific cCREs, their target genes, and the transcription factors (TFs) that may mediate their effects on gene expression changes in LOAD. We found that the expression of a subset of these TFs also changed in LOAD in the same specific cell subtype. Moreover, focusing on LOAD-GWAS regions, we catalogued putative regulatory SNPs positioned in the identified LOAD-associated cell subtype-specific cCREs that change the affinities of TF motifs and thereby potentially affect the expression of the target genes. Collectively, we provided new insights into the relationships between DNA sequence variation, chromatin structure, and transcriptome in LOAD brains at an unprecedented cell subtype-specific resolution. Furthermore, we identified candidate *cis-* and *trans-* regulatory networks and their target genes for further experimental investigations in model systems relevant to LOAD.

**Figure 1.**
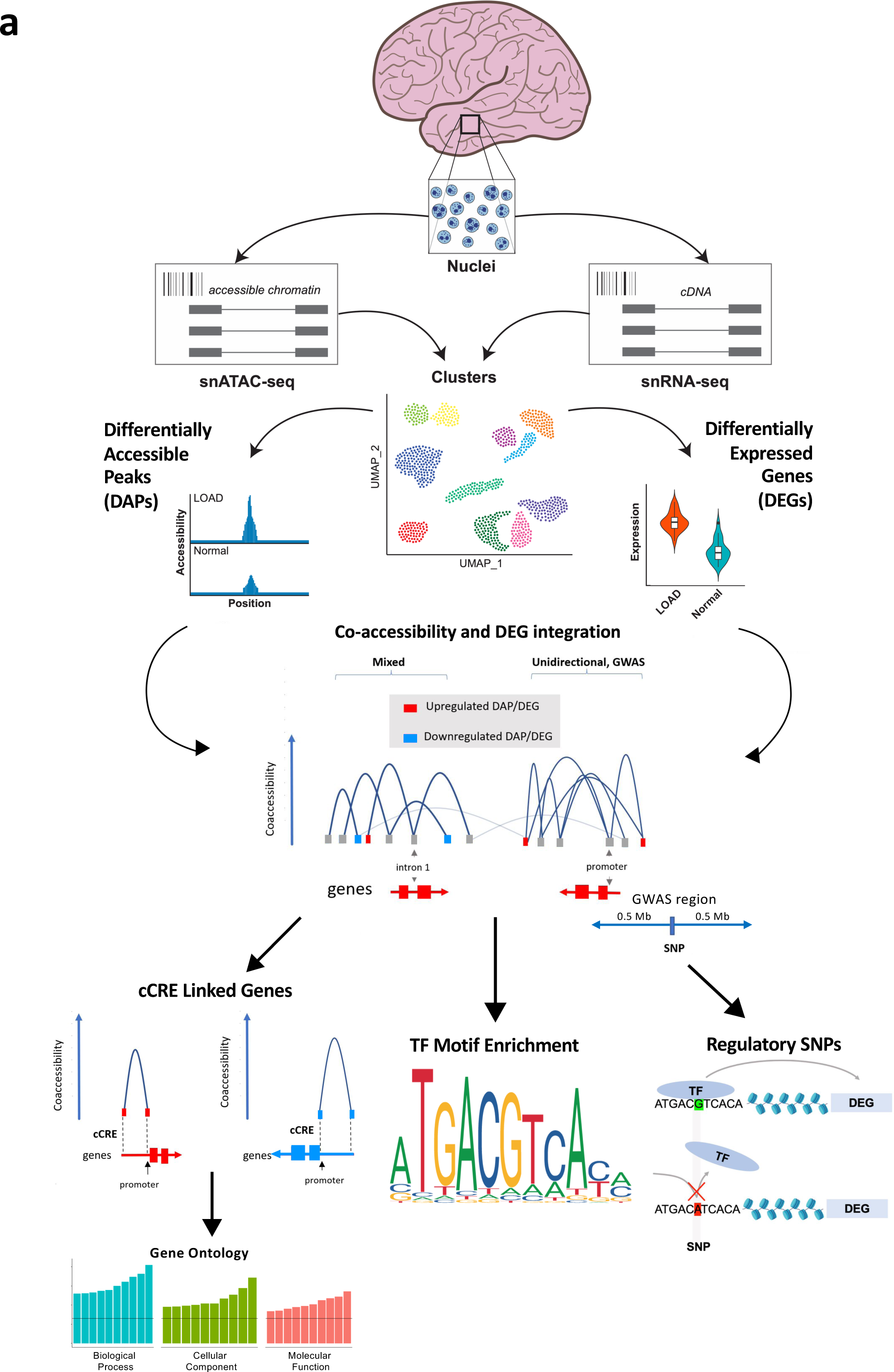
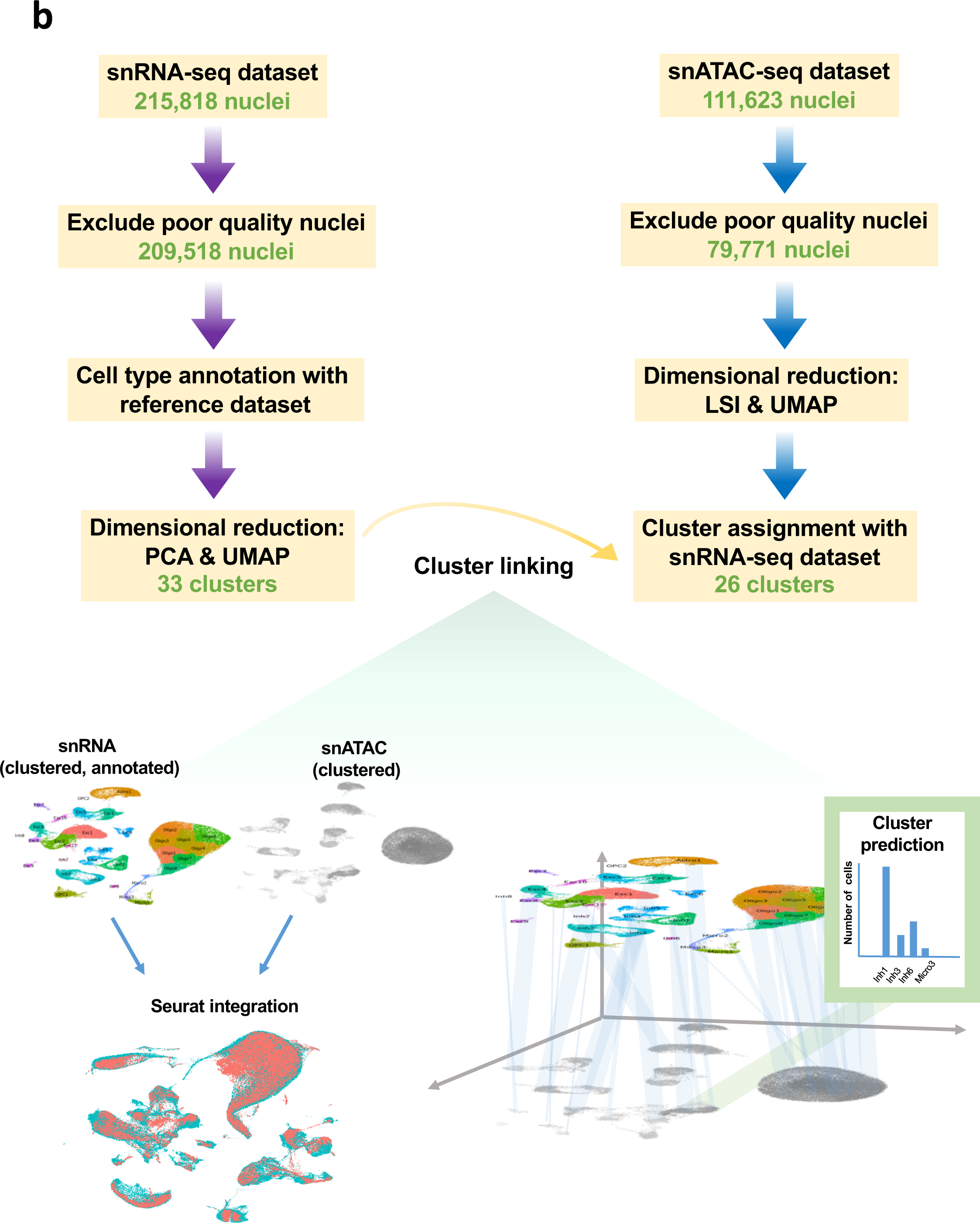
Experimental approach and integration of snATAC-seq with snRNA-seq clusters. **a**, Schematic of nuclei isolation and parallel snATAC-seq and snRNA-seq library generation. Tissue ➔ nuclei ➔ parallel library generation ➔ matching clusters ➔ DEGs/DAPs ➔ coaccessibility ➔ 1Mb DEGs, enriched TFs. **b**, Flow chart of snRNA-seq and snATAC-seq analytical pipeline leading to cluster assignment, and schematic of analytical strategy with label transfer of snRNA-seq clusters onto snATAC-seq data by cell type and subtype.

## RESULTS

### Characterization of cell types and subtypes in the human temporal cortex of healthy aging and Alzheimer’s individuals using multi-omics datasets

We isolated nuclei samples from frozen post-mortem human temporal cortex (TC) tissues of 12 LOAD and 12 cognitively normal individuals (Table 1, Table S1) and performed both snRNA-seq and snATAC-seq in parallel by simultaneously using two aliquots from the same nuclei sample (Fig. 1a). To exclude the APOEe4 effect, all donors were APOEe3/3. In both snRNA-seq and snATAC-seq nuclei, we discovered multiple neuronal and glial cell subtypes which were linked across both data sets for downstream analyses (Fig 1b). This parallel experimental design allowed us to minimize differences between the snRNA-seq and snATAC-seq assays, including technical variables and variability in cell type composition, and facilitated the multi-omics integrative analyses outlined in Fig 1.

**Table 1.**
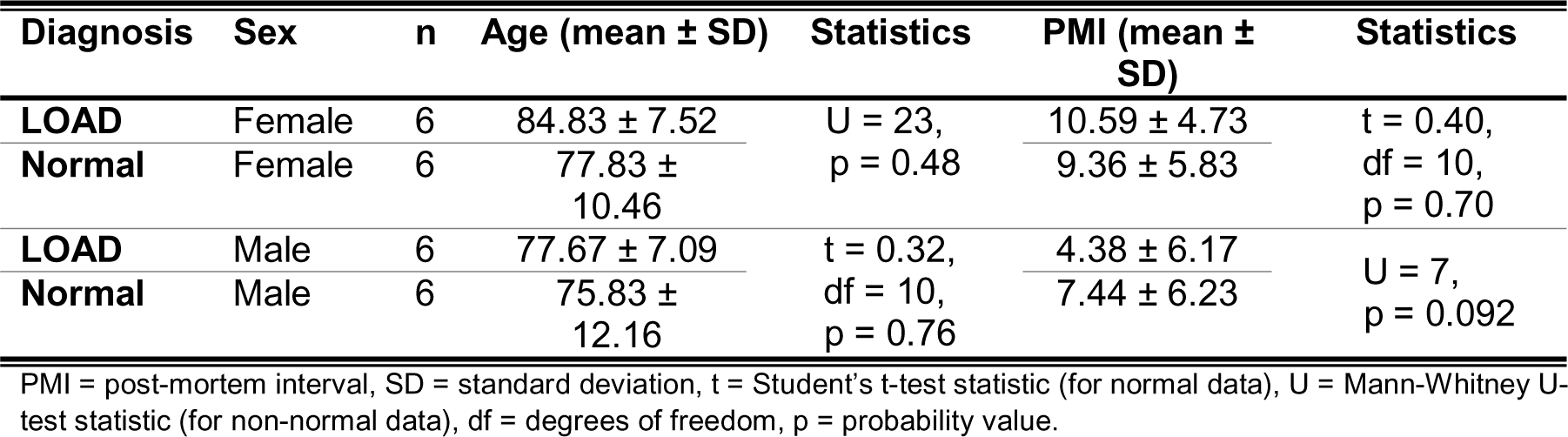
Demographics summary of study cohort.

We first annotated the cell types in our snRNA-seq dataset by the label transfer method^38^ using a pre-annotated reference snRNA-seq dataset generated with the same technology^33^, and validated these annotations using known cell type-specific markers (Methods, Fig. S1a). After quality control (QC) filtering, we retained a total of 209,518 nuclei for snRNA-seq from all 24 temporal cortex samples (Table S2). After dimensionality reduction followed by Louvain community detection, we identified 33 distinct cell clusters (Figs. 2a, 3a, Tables S2-4). Clusters of nuclei representing cell subtypes were each given a unique label according to their broader cell type (*e.g.,* the 11 clusters of excitatory neurons were labeled Exc1-Exc11, Table S4). Previously known neuronal subtypes such as somatostatin interneurons^39–41^ tended to form separate clusters and a particular subtype of excitatory neurons marked by *LAMP5* was notably depleted in LOAD tissue (Fig. S1b-c). Separate clustering analysis for normal and LOAD nuclei samples did not yield qualitatively distinct patterns of clusters between the two datasets, suggesting the presence of broadly similar cell subtypes regardless of disease status (Fig. S2).

**Figure 2.**
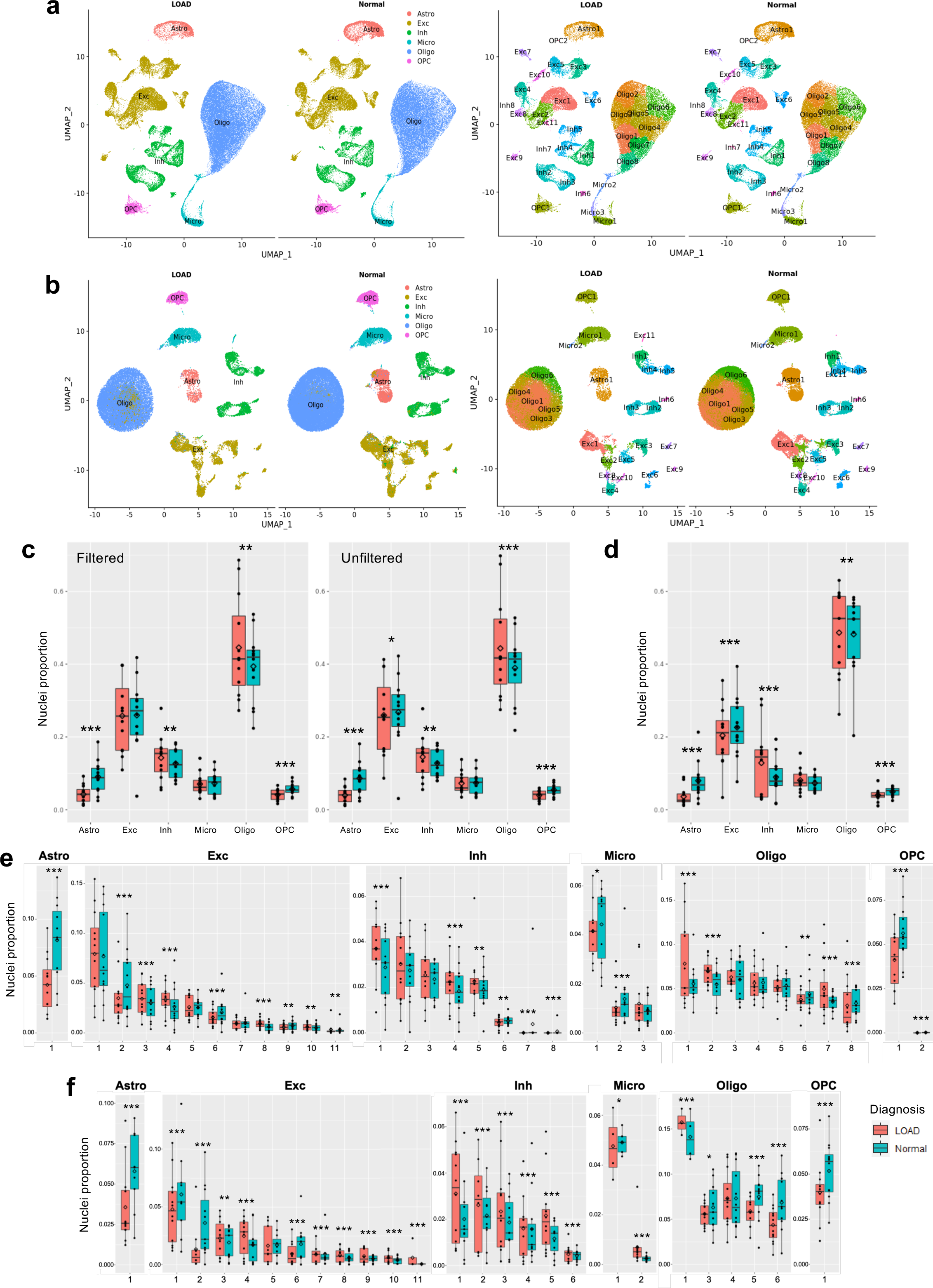
Proportions of LOAD and normal nuclei among cell types and subtype clusters. **a** and **b**, Uniform manifold approximation and projection (UMAP) dimensional reduction plots of (a) snRNA-seq and (b) snRNA-seq datasets split into LOAD and normal groups following integration and clustering of the total nuclei population. Cell types are labeled in left-hand plots and cell type subclusters are shown in right-hand plots. **c**, Box plots quantifying differences in cell type proportions between LOAD and normal samples based on snRNA-seq dataset. As indicated, plots represent cell datasets following QC filtering and without QC filtering. Dots represent means for each sample set. Boxes represent interquartile range, with median values indicated via horizontal line. Mean values are indicated by open diamonds. Whiskers extend to sample means within 1.5 times the interquartile range. Asterisks represent statistical significance of variation between LOAD and normal mean values at levels of *p* < 0.05 (*), *p* < 0.01 (**) and *p* < 0.001 (***) based on bootstrapped Wilcoxon test. **d**, Box plots quantifying differences in cell type proportions between LOAD and normal samples based on snATAC-seq dataset. **e** and **f**, Box plots quantifying differences in subcluster proportions between LOAD and normal samples based on (e) snRNA-seq and (f) snATAC-seq datasets.

**Figure 3.**
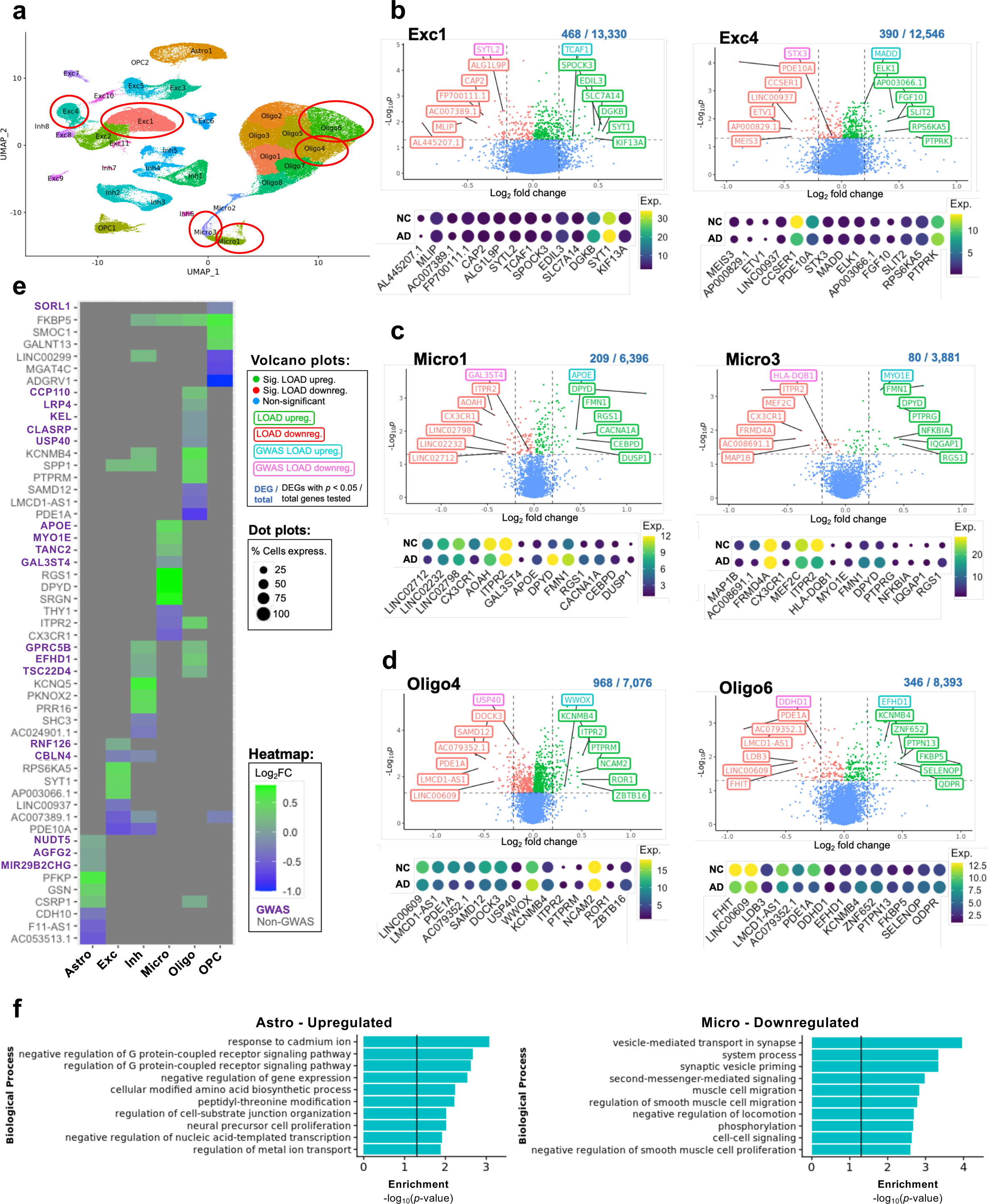
Top differentially-expressed genes (DEGs) upregulated and down-regulated in LOAD by cluster and cell type. **a,** UMAP dimensional reduction plot of 33 cell subtype clusters based on snRNA-seq data. Clusters highlighted in panels b-d are circled in red. **b-d**, Unbiased volcano plots for six example clusters representing excitatory neuron (Exc), microglia (Micro), and oligodendrocyte (Oligo) cell types. Log_2_ fold change (FC) between LOAD and normal control (NC) samples is plotted against –log_10_ *p*-value (FDR). Points representing DEGs with statistically significant (*p* < 0.05) upregulation in LOAD are shown in green while DEGs with significant downregulation are shown in red. Genes without significantly differential expression are shown in blue. The proportion of DEGs to total genes examined is shown above each plot. The six DEGs with the highest absolute fold change (log_2_ FC > 0.2) in the up- and downregulated categories are labeled in green and red, respectively. The top up- and downregulated DEGs within 500kb of disease-associated SNPs previously identified in GWAS are labeled in teal and pink, respectively. For each of the labeled genes, dot plots are shown below representing their unscaled expression levels (color) and percent of cells expressing the gene (width) for LOAD and NC samples. **e**, Heatmap showing log_2_ FC for the top three DEGs with the highest fold changes in upregulated (green) and down-regulated (blue) categories for each cell type examined, as well as the top three DEGs within 500kb of identified GWAS SNPs (labeled in purple). **f**, The top ten GO terms for upregulated LOAD genes (all significant, FDR adjusted *p*-values) for astrocyte and downregulated LOAD genes for microglial cell types.

Similarly, 79,771 nuclei for snATAC-seq were retained after QC filtering and Louvain community detection yielded 26 cell subtype clusters (Fig. 2b, 4a, Table S5). We mapped the resulting snATAC-seq clusters to the corresponding snRNA-seq clusters through the identification of shared transfer anchors between the datasets based on observed gene expression profiles for snRNA-seq nuclei and predicted gene expression profiles for snATAC-seq nuclei (Fig. 1b). snATAC-seq clusters were linked to most snRNA-seq clusters with the exception of seven cell subtypes: inhibitory neurons (Inh)7, Inh8, microglia (Micro)3, oligodendrocytes (Oligo)2, Oligo7, Oligo8, and oligodendrocyte precursor cells (OPC)2. Validation of the cell subtypes linking accuracy showed an overall 75% accuracy rate (Jaccard index) and high concordance with a mean of ∼0.8 (hybrid score)^35^ (Table S6).

LOAD brains are pathologically characterized by neuronal loss and gliosis^42^. Thus, we examined differences in cell type and subtype proportions between LOAD and normal samples. Using the snRNA-seq dataset from the quality control (QC)-filtered nuclei we observed significant decreases in the proportions of astrocyte and OPC nuclei (*p* < 0.001) and increases in the proportions of inhibitory neurons and oligodendrocytes nuclei (*p* < 0.01) in the LOAD samples (Fig. 2c, Tables S2 and S4). Extending the analysis to the full nuclei set, without removing the data from lower quality nuclei (based on QC parameters), replicated the same observations and additionally showed a decrease in the proportion of excitatory neurons (*p* < 0.05, Fig. 2c), as expected in LOAD brains. Supporting these findings, all significant changes in the proportions of LOAD nuclei compared to controls were also detected in QC-filtered snATAC-seq nuclei (*p* < 0.01) (Fig. 2d, Table S5).

We then performed the nuclei proportion comparison on more granular level of cell subtypes. For the majority of the cell subtypes, the snRNA-seq and snATAC-seq QC-filtered nuclei demonstrated differences in proportion between LOAD and control, and the directionality of most differences were consistent between these datasets (Fig. 2e and f, and Tables S2 and S5, respectively). However, we observed some discrepancies in directionality between the snRNA-seq and the snATAC-seq datasets for the smaller subclusters comprising less than 2% of total nuclei (e.g., Exc9, Inh6, Micro2), where even small deviations would be expected to have a greater impact on observed proportions. Several subclusters within particular cell-types showed the same directions of proportional differences between LOAD and control as those reported previously^36^. Furthermore, similarly to the previous observations^36^, we found that within the same cell-type some subclusters were significantly enriched in LOAD, whereas others were depleted (Fig 2e, f). Nonetheless, the overall proportions of cell types and subtypes may not reflect true proportions in the brain due to some cell types not being equally amenable to the nuclei preparation procedure.

### Cell type- and subtype-specific differential gene expression and chromatin accessibility in LOAD

We used the snRNA-seq and snATAC-seq datasets to characterize LOAD-associated dysregulated genes and changes in chromatin structure within each cell type and subtype. To examine differential gene expression using the snRNA-seq data, we utilized a mixed effects model that incorporates a random effect for donor to avoid pseudo-replication bias^43^ (Methods), which is typically not accounted for in single-cell studies. In addition to diagnosis, age, sex, post-mortem interval (PMI), sequencing saturation, and nuclei proportion within each cluster were incorporated into the model as fixed effects, based on our covariate analysis of 38 metadata variables (Methods, Fig. S3, Table S2).

This analysis identified numerous differentially expressed genes (DEGs) in LOAD for each cell type (ranging from 135 in OPCs to 429 in oligodendrocytes) and subtype (ranging from zero in subtype clusters Exc11, Inh7 and 8, and OPC2, to 968 in cluster Oligo4, Tables S7 and S8). The DEGs that showed the strongest (based on |log_2_FC|) and most significant (false discovery rate (FDR)-adjusted *p*-value) upregulation and downregulation effects in LOAD are highlighted in Fig. 3a-d and Fig. S4. In addition, we indicated the top DEGs that are within 500kb (upstream or downstream) of a LOAD GWAS tag SNP ^8^ (Fig. 3a-d, Fig. S4). Some of these DEGs were the most proximate to the tag SNP (e.g., *APOE* in Micro1 and *WWOX* in Oligo4), however, many were more distal within the tag SNP ±500kb window (Fig. 3c, d). For example, while *EPHA1* is the nearest gene to LOAD-GWA SNP rs10808026, *TCAF1* was identified as a DEG in Exc1 within this LOAD region (Fig. 3b).

The differential expression analysis at the cell type level found that the majority of the strongest DEGs did not co-occur across cell types, supporting the need for cell type specificity in genomic analyses of LOAD. The common DEGs showed more subtle changes, and few had opposite directionality between cell types (Fig. 3e).

We used the upregulated and downregulated DEGs in LOAD from each cell type to perform gene ontology (GO) analysis, revealing enrichment of biological pathways relevant to neuronal function and neurological disease processes in several cell types (Fig. 3f and Fig. S3). For example, the GO term ‘vesicle mediated transport in synapse’ was enriched for downregulated genes in LOAD for microglia and ‘response to cadmium ion’ (involved in neurotoxicity, oxidative stress, and neuronal apoptosis) for upregulated genes in LOAD for astrocytes (Fig. 3f).

Using snATAC-seq, we profiled accessible chromatin regions (peaks) and catalogued cell type and subtype-specific differentially accessible peaks (DAPs). A differential analysis approach revealed many DAPs between LOAD and normal for each cell type (ranging from 25 in astrocytes to 44,703 in excitatory neurons) and subtype (ranging from zero in clusters Exc11, Inh7 and 8, Micro3, Oligo2, 7 and 8, and OPC2, to 53,661 in Exc1). We thus identified LOAD-associated DAPs at both cell type and cell subtype resolutions (Fig 4 a-c, Tables S7 and S8). At the cell subtype level, we showed four representative examples, and for each we indicated the top DAPs with the strongest (based on |log_2_FC|) and most significant (FDR-adjusted *p*-value) LOAD-associated effects on chromatin accessibility, as well as the top DAPs within GWAS loci^7^ (tag SNP±500kb; Fig. 4a,b). We also show the top DAPs for all cell types and highlighted those that overlapped with GWAS loci (Fig. 4c). DAPs were labeled by distance to the nearest gene. Several DAPs, such as *SCAF11,* were identified in a few cell types, while others were unique for a specific cell type, such as the DAPs near the *PTK2B* and *INPP5D* genes in microglia (Fig.4c).

**Figure 4.**
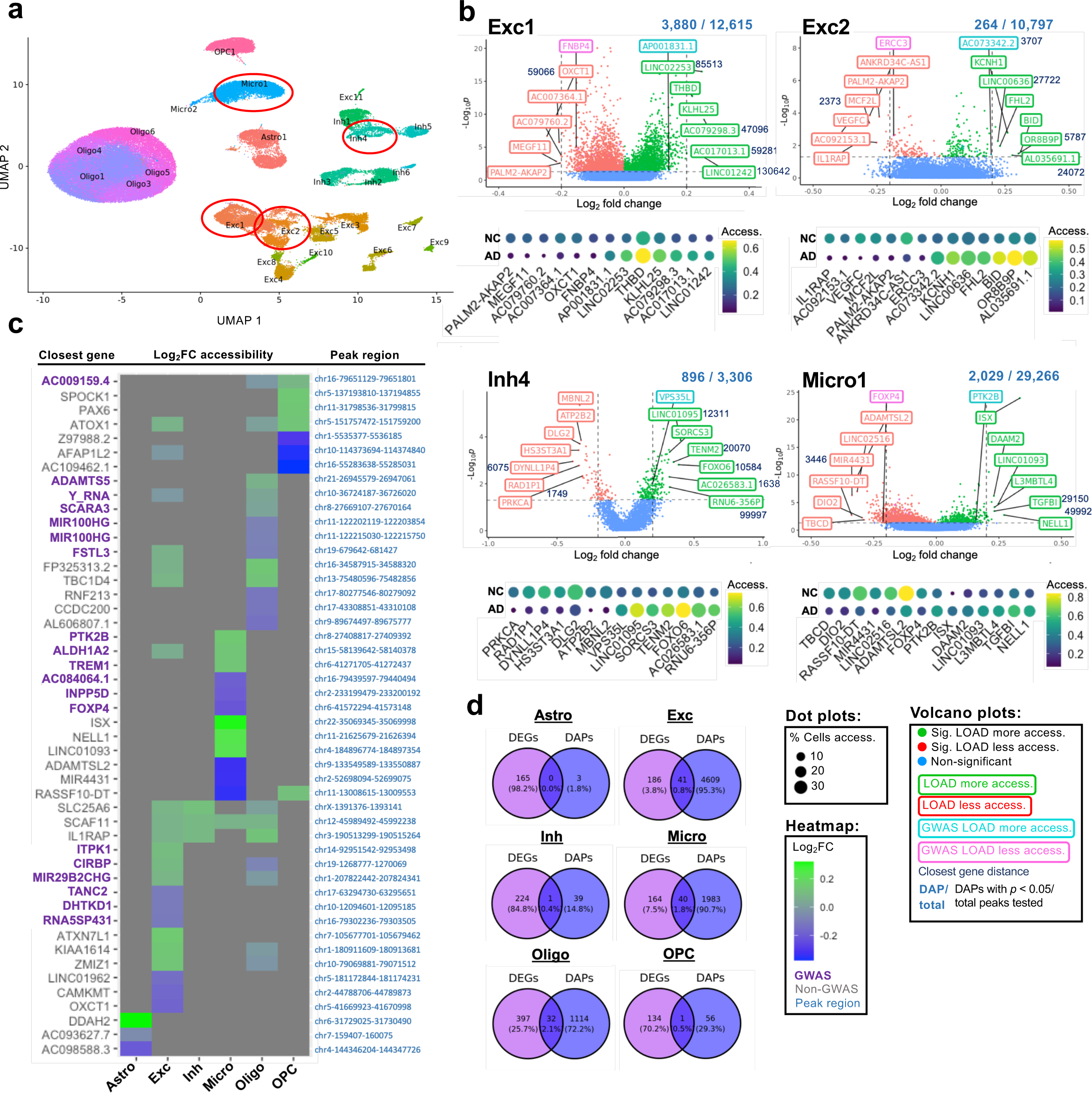
Top differentially-accessible peaks (DAPs) upregulated and down-regulated in LOAD by cluster and cell type. **a**, UMAP dimensional reduction plot of snATAC-seq cell data indicating both cell type and subtype clusters. Clusters highlighted in panel b are circled in red. **b**, Unbiased volcano plots for four example clusters representing excitatory neuron, inhibitory neuron, and microglia cell types. Log_2_ fold change (FC) between LOAD and normal control (NC) samples is plotted against –log_10_ *p*-value (FDR). Points representing DAPs with statistically significant (*p* < 0.05) upregulation of accessibility in LOAD are shown in green while DAPs with significant downregulation are shown in red. Peaks without significantly differential accessibility are shown in blue. The proportion of DAPs to total peaks examined is shown above each plot. The closest genes to the six DAPs with the highest absolute fold change (log_2_ FC > 0.2) in the up- and downregulated categories are labeled in green and red, respectively. The closest genes to the top up- and downregulated DAPs within 500kb of GWAS SNPs are labeled in teal and pink, respectively. Distances of closest genes from corresponding peaks (in bp) are indicated adjacent to gene labels where such distances are greater than zero. For each of the labeled peaks, dot plots are shown below representing their unscaled accessibility levels (color) and percent of cells with identified peak accessibility (width) for LOAD and NC samples. **c**, Heatmap showing log_2_ FC for the top three DAPs with the highest fold changes in upregulated (green) and down-regulated (blue) categories for each cell type examined, as well as the top three DAPs within 500kb of identified GWAS SNPs (labeled in purple). Closest gene names are indicated to the left of the heatmap while peak ranges are shown to the right. **d**, Venn diagrams showing overlap between DEGs identified via snRNA-seq and DAPs identified via snATAC-seq for each cell type examined.

### *Cis* co-accessibility network analysis in LOAD identified cell subtype specific DEGs linked to cCREs

Next, we examined the crosstalk between cCREs and gene expression in LOAD TC. We defined a cCRE as a noncoding DAP that is co-accessible with one or more peaks overlapping the promoter/intron1 of a DEG. Comparative analysis between the DEGs and DAPs closest genes for each cell type found a relatively small degree (0.4-2.1%) of overlap between the two gene sets (Fig. 3d), possibly due, at least in part, to the challenges in accurately assigning DAPs to target genes. These results suggest a complex interaction between LOAD-associated cCREs and target genes, and that cCREs may not simply regulate their nearest genes.

To address this complexity, we performed a multi omics integrative approach using our *parallel* snRNA-seq and snATAC-seq datasets to characterize the *cis*-regulatory landscape in LOAD and to identify target genes of LOAD-associated cCREs. To this end we used the snATAC-seq data from the LOAD nuclei only and utilized the Cicero algorithm^44^ (Methods) to construct *cis*-co-accessibility networks (CCANs) for each cell subtype. We defined a total of 25,020 CCANs in LOAD TC across all 26 snATAC-seq cell subtypes and provided a detailed summary of their characteristics (Table S9). To identify CCANs with likely specific relevance to LOAD, we further refined this set to include only CCANs that contained at least one LOAD-associated DAP and DEG identified in our differential analyses of the snATAC-seq and snRNA-seq datasets, respectively (described above). This filtering criteria identified a subset of 518 LOAD CCANs in 15 cell subtypes that were categorized into 3 groups based on the directionality (log_2_FC) of the DAP-DEG pair: (i) unidirectional (309 CCANs), all more accessible/upregulated or less accessible/downregulated; (ii) mixed (129 CCANs), both directions for the DAPs and/or DEGs whereas at least one DAP-DEG pair showed the same effect direction; (iii) bidirectional (80 CCANs), all opposite directions. (Fig 1a, Fig. 5a-c, Table S9). Further analyses focused on the unidirectional and mixed LOAD CCANs categories (Fig 5), several of which (4 and 3, respectively, across Exc1, Micro1, Oligo4 and Oligo6 cell subtypes) also overlapped with LOAD-GWAS loci^7^. To identify the target genes of LOAD-associated cCREs in each cell subtype, we examined the co-accessibility of the peak overlapping the DEG promoter or intron 1 region with the distal DAP that defined the LOAD-associated cCRE. We catalogued the cCRE-linked genes for which their promoter/intron 1 peak was also a DAP and showed the same directionality as the cCRE-DAP. The analysis revealed 69 DEGs linked to cCRE in 8 of the cell subtypes with expression fold change (log_2_FC) greater than 0.15 (Fig 5d, Tables S9 and S10). 32 of the target DEGs were linked to 2-11 cCREs including, *APOE* and *BIN1*, regulated by 3 and 2 cCREs, respectively (Fig. 5d, e, Tables S9 and S10). On the other hand, no LOAD cCREs were linked to more than one DEG (Table S10). A number of the identified cCRE-targeted DEGs have been implicated in neurodegenerative diseases, including genes involved in AD pathogenesis such as, *MAPK3*, *APP*, and *FAM107B* in excitatory neuron cluster 1 (Exc1), *MYO1E* and *APOE* in microglia cluster 1 (Micro1), and *BIN1* in oligodendrocyte cluster 4 (Oligo4) (Fig. 5d, Table S9). Collectively, our new strategy led to the identification of novel LOAD genes as well as validation of known disease genes and suggested the cell subtype in which they exert their pathogenic effects. In addition, we characterized the *cis*-regulatory elements and networks that govern the dysregulation of these genes in disease.

**Figure 5.**
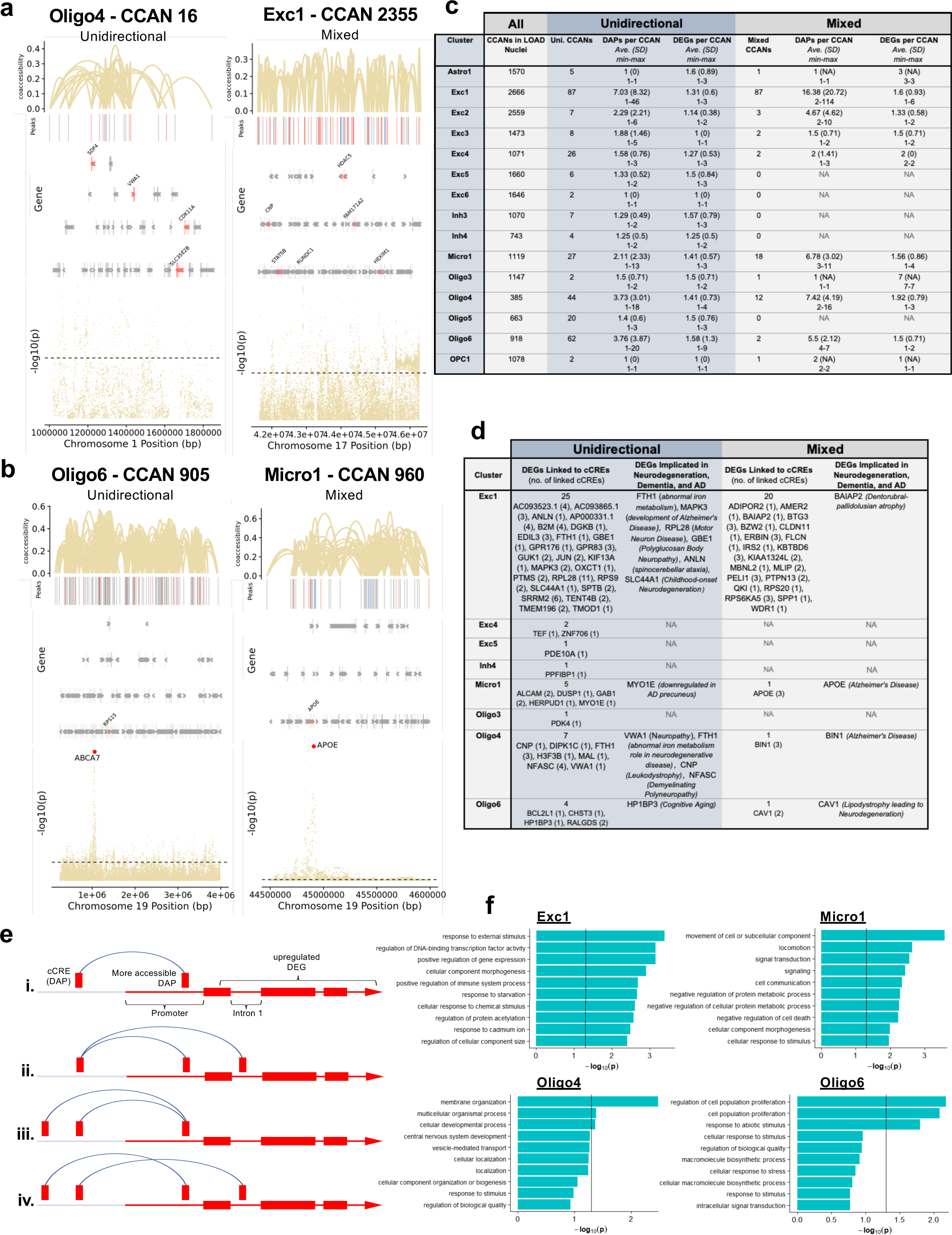
Identification of cis co-accessible networks and associated DEGs in LOAD nuclei. **a** and **b**, diagrams of example unidirectional CCANs - in which all overlapping DAPs and DEGs are regulated in the same direction in LOAD nuclei (e.g. more accessible peaks and higher gene expression) and mixed CCANs - in which at least one DAP/DEG pair are regulated in the same direction and at least one pair are regulated in opposite directions. Vertical lines at the top of each diagram indicate chromatin peaks and regulatory linkages of accessibility between peak pairs and associated co-accessibility scores are indicated by Bezier curves. Only CCAN-associated peaks are shown. DAPs with greater accessibility in LOAD are shown in salmon, while those with reduced accessibility in LOAD are shown in blue. Non-differentially-accessible peaks are shown in grey. Below the peak linkage plots, gene exons overlapping CCAN regions are depicted as arrows indicating the directionality of transcription. DEGs are labeled with gene names, and LOAD-upregulated DEGs are shown in salmon while non-differentially expressed genes are shown in grey (no downregulated DEGs depicted). At the bottom of each diagram, a Manhattan plot is shown indicating -log_10_ of *p*-values for LOAD-association of chromosome loci within CCAN region as calculated by Kunkle *et al*.^7^ Dotted lines in Manhattan plots indicate statistical significance threshold (*p* = 0.05). Panel a depicts example CCANs that do not overlap GWAS-identified LOAD-associated SNPs, while panel b depicts example CCANs that do overlap LOAD-associated SNP loci, indicated via orange dots and labeled on Manhattan plots. **c**, Table showing total number of CCANs identified for each snATAC-seq nuclei cluster, as well as the number of unidirectional and mixed CCANs, and the mean number of DAPs and DEGs identified per CCAN for each cluster in both the unidirectional and mixed categories, along with the standard deviation (SD) and maximum and minimum values. **d**, Table listing DEGs linked to cCREs in unidirectional and mixed CCANs for each applicable cluster, as well a functional annotation of specific DEGs with known associations to neurodegenerative disease. Number of cCREs linked to each DEG is shown in parentheses. **e**, Diagram of potential DEG/cCRE associations in which i.) one DAP is coaccessible with one peak overlapping the DEG, ii.) one DAP is coaccessible with multiple DEG peaks, iii.) two DAPs are coaccessible with one DEG peak, and iv.) two DAPs are coaccessible with two DEG peaks. **f**, Gene ontological analysis of biological processes for cCRE-linked DEGs associated with CCANs in the indicated cell subtype clusters. The top ten enriched biological processes involving a minimum of three DEGs are listed.

GO analysis of DEGs targeted by cCREs in each cell subtype revealed a few enriched biological processes (Fig. 5f, Fig. S6). For example, Exc1 showed enrichment for GO terms related to gene transcriptional regulation and response to external stressors. In Micro1, cCRE-linked DEGs involved in biological processes related to signal transduction, metabolic process and movement of cellular components were among the most enriched. Oligo4 and Oligo6 showed enrichment for GO terms involved in neural cell development, and cell proliferation, respectively. Due to the relatively small numbers of genes identified as LOAD associated cCRE-targeted DEGs specific to these cell subtypes, enrichment scores for the reported GO terms were not statistically significant but still suggested trends.

### Cell subtype specific transcription factors relevant to LOAD

We sought to explore the cell subtype specific *trans* regulation involved in LOAD-associated changes in gene expression as a complementary approach to our *cis*-regulation analysis. To this end, we searched transcription factors (TFs) that may interact with LOAD CCANs and thus contribute to dysregulation of genes in LOAD. The analysis focused on the subset of unidirectional and mixed LOAD CCANs identified in each of the 15 cell subtypes as described above (Fig. 5c) and was inclusive to all peaks (all accessible chromatin sequences, not limited to DAPs). The HOMER software package was used to identify enrichment of TF binding sites within the subset of LOAD CCANs for each cell subtype for TFs that were expressed in ≥10% of the corresponding cell subtype (Fig. 6a, Table S11). The analysis revealed enrichment for 17 to 124 TFs in 7 out of the 15 analyzed cell subtypes. Interestingly, we found that several of these TFs were also LOAD-associated DEGs in five cell subtypes including, Exc1, Exc4, Micro1, Oligo4, and Oligo5 (Fig. 6a-c, Table S11). *ELK1* - enriched in Exc1 - showed the highest fold increase in expression in LOAD vs. normal cells (log_2_FC = 1.32, Fig. 6b-c). *ELK1* has been previously shown to initiate regionalized neuronal death and to associate with inclusions present in Alzheimer’s disease, Lewy body disease, and Huntington’s disease^45^. Other TF examples included *ATF7*, involved in stress-responsive and heterochromatin formation and previously identified as a probable LOAD-susceptibility marker^46^, showed the highest enrichment in Exc1 and Oligo5; *KLF13*, a key regulator of schizophrenia^47^, was enriched in Micro1 and Oligo4; *JUN* and its heterodimers with members of the *FOS* family^48^, were enriched in Exc1 (Fig. 6b-c). Altogether, these findings point to key TFs, regulatory elements, and *cis-trans* interactions potentially involved in dysregulating gene networks in specific cell subtypes contributing to LOAD pathogenesis.

**Figure 6.**
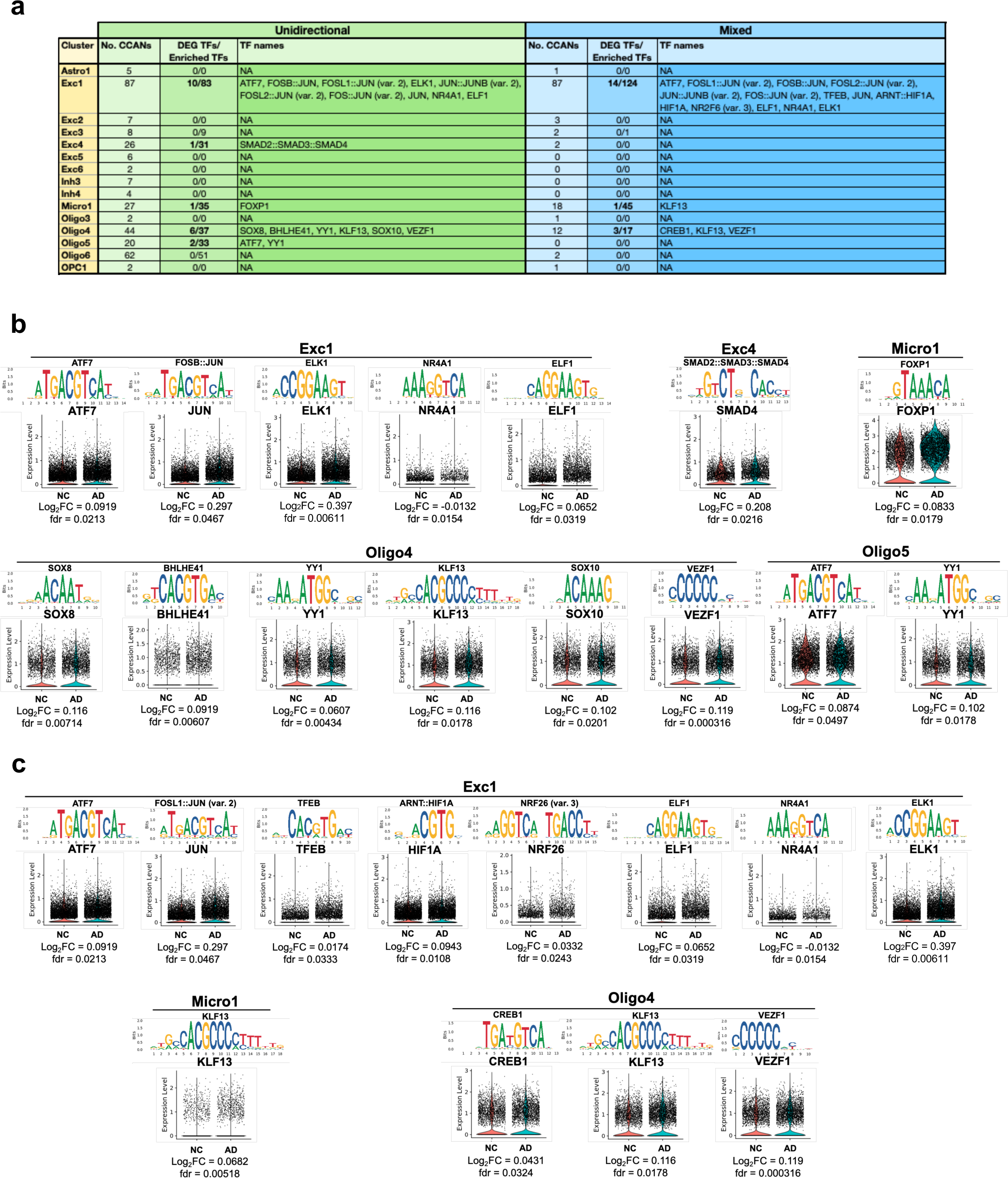
Differential expression of transcription factors with motifs enriched in cis co-accessible networks. **a,** Table showing number of CCANs identified for each snATAC-seq cluster as well as the number of enriched transcription factor (TF) motifs corresponding to TFs identified as DEGs in our snRNA-seq analysis within both unidirectional and mixed CCANs. The specific identified TFs are indicated where applicable. **b** and **c**, Sequence logos for indicated enriched motifs in unidirectional (b) and mixed (c) CCANs associated with labeled DEG TFs, along with violin plots of normalized snRNA-seq expression data split by NC and LOAD groups for the corresponding TFs for each applicable cluster. Also indicated are *p*-values (FDR) and log_2_FC for TF gene expression data. Motifs are presented from left to right in decreasing order of enrichment for each cluster. Where multiple motifs corresponded to the same TF, the most highly enriched motif is shown.

### Identification of SNPs potentially affecting TF binding and expression of DEGs in LOAD-GWAS regions

GWAS have previously identified numerous LOAD-associated SNPs, termed tagging SNPs. However, the majority of these tagging SNPs are based on disease association only, and the actual variants involved in disease risk have yet to be identified. We sought to prioritize candidate functional SNPs within GWAS regions that directly affect LOAD risk, specifically via transcriptional mechanisms. To this end we performed the analysis on 21 LOAD CCANs (unidirectional and mixed) that also overlapped LOAD-GWAS regions (Fig. 7a, Table S12). We further focused the analysis on SNPs overlapping predicted TF binding sites for the minor or the major allele (p<1×10^-5^) with MAF≥1%. Next, we catalogued 125 SNPs that were predicted to change the affinity of 303 TF binding motifs (FDR<0.05) using the Affinity Test for Identifying Regulatory SNPs (atSNP) R software package^49^ (Fig. 7a, Fig. S7a, Table S12). The minor alleles of the identified candidate regulatory SNPs resulted in gain or loss of TF binding sites in a cell subtype-specific manner. These identified candidate regulatory SNPs mapped within LOAD-associated cCREs sequences linked to 16 DEGs across 4 cell subtypes (Exc1, Micro1, Oligo4 and 6, Fig. 7a, Table S12). In the majority of the cCRE linked-DEGs (11 of 16), the peak overlapping the DEG promoter/intron 1 was also a DAP (Table S12, examples in Fig. 7b-e and Fig. S7). These candidate regulatory SNPs were identified in specific cell subtypes. Many of the SNPs were identified in Exc1 (Fig. 7a, Table S12), and presumably affect expression of genes including *CUTA*, a negative regulator of beta-amyloid generation ^50^, *TFEB*, a TF regulator of autophagic dysfunction associated with neurodegenerative pathology ^51^, *FZR1,* an adapter protein involved in cell cycle regulation that may suppress Cyclin B levels affecting AD-associated aberrant cell cycle re-entry ^52^, *GNA11*, associated with hypocaliciuric hypercalcemia type II ^53^, *RPS15*, a structural ribosome component linked to Parkinson’s disease ^54^ (Fig. 7b, Fig. S7a-b, Tables S12 and S13), and *MBD3,* a neuropathy-associated chromatin remodeling complex component^55^. In Micro1, DEGs affected by the catalogued candidate regulatory SNPs included *MYO1E*, an actin-based molecular motor protein previously found to be differentially expressed in a microglial model of AD ^56, 57^ (Fig.7c, Fig. S7c-f, Tables S12 and S13) and *APOE* ^7, 8^ (Fig. 7d, Fig. S7g-i, Tables S12 and S13). In Oligo4 we identified SNPs affecting *BIN1,* a nucleocytoplasmic adaptor protein involved in synaptic vesicle endocytosis in the CNS (Fig. 7e, Fig. S7j-l, Tables S12 and S13). *BIN1*, a major risk for LOAD, was suggested to have a role in regulating postsynaptic trafficking, and to accelerate beta-amyloid levels and tau accumulations in the context of LOAD ^58–61^. Other DEGs affected by the identified candidate SNPs in Oligo4 included *NDUFS7*, a mitochondrial respiratory complex component associated with several neurological disorders^62^, *PTBP1*, a regulator of neuronal pre-mRNA splicing associated with frontotemporal dementia and amyotrophic lateral sclerosis^63, 64^ and SC5D a cholesterol biosynthesis enzyme associated with lathosterolosis, a congenital disorder affecting central nervous system development ^65^, and *RPS15* ^54^ in Oligo6 (Table S12). Overall, many DEGs for which expression was predicted to be influenced by these identified SNPs were implicated in LOAD and other neurodegenerative disorders. Only 4 DEGs did not have previous evidence related to neurological phenotypes, *EFHD1*, *TAPBP* and *TCAF1* in Exc1, and *LST1* in Micro1. These findings support the biological relevance of this methodology in identifying cell subtype-specific variants with putative roles in LOAD pathogenesis. Noteworthily, many of the candidate regulatory SNPs changed the binding affinities of several TF motifs and each of the 16 cCRE linked-DEGs was associated with multiple SNP-TF interactions (Fig. 7a, Table S12), however, some of these TFs may not be of biological relevance. Thus, the subsequent analysis was restricted to TFs that were expressed in ≥10% of the corresponding cell subtype. In addition, to prioritize the top candidate regulatory SNPs we considered the affected genes and therefore narrowed down to cCRE linked-DEGs with |log_2_FC|≥0.15. After applying these criteria, the analysis revealed a total of 20 candidate regulatory SNPs and 5 candidate dysregulated genes including 10 SNPs in Micro1 presumably exert their effects on *MYO1E* or *APOE*, 7 SNPs in Oligo4 potentially affecting *BIN1*, and 3 SNPs for in Exc1 *RPS15* or *TCAF1* (Fig. 7b-e, Fig. S7 b-m, Table S13). Collectively, these outcomes provided a high priority list of candidate regulatory SNPs, TFs and target genes for experimental validation studies using model systems that facilitate investigations in the specific relevant brain cell subtype.

**Figure 7.**
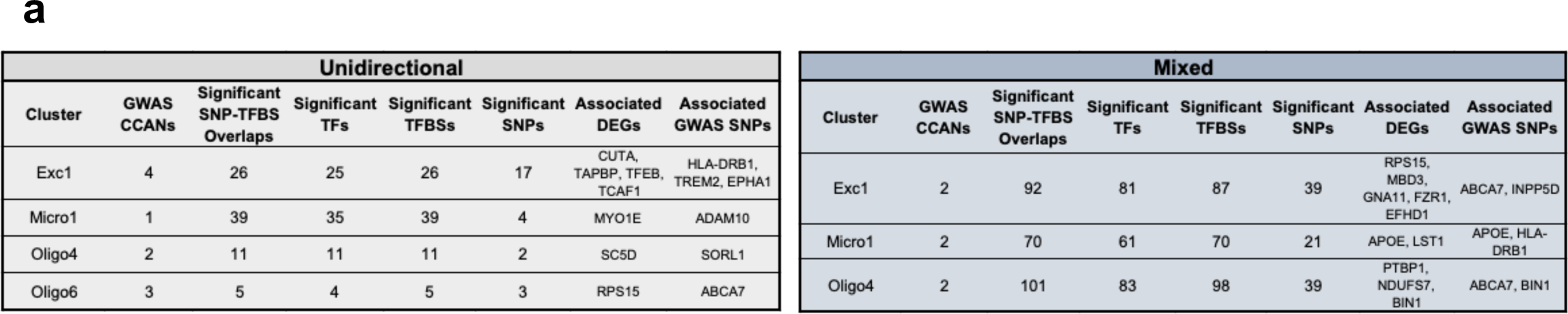
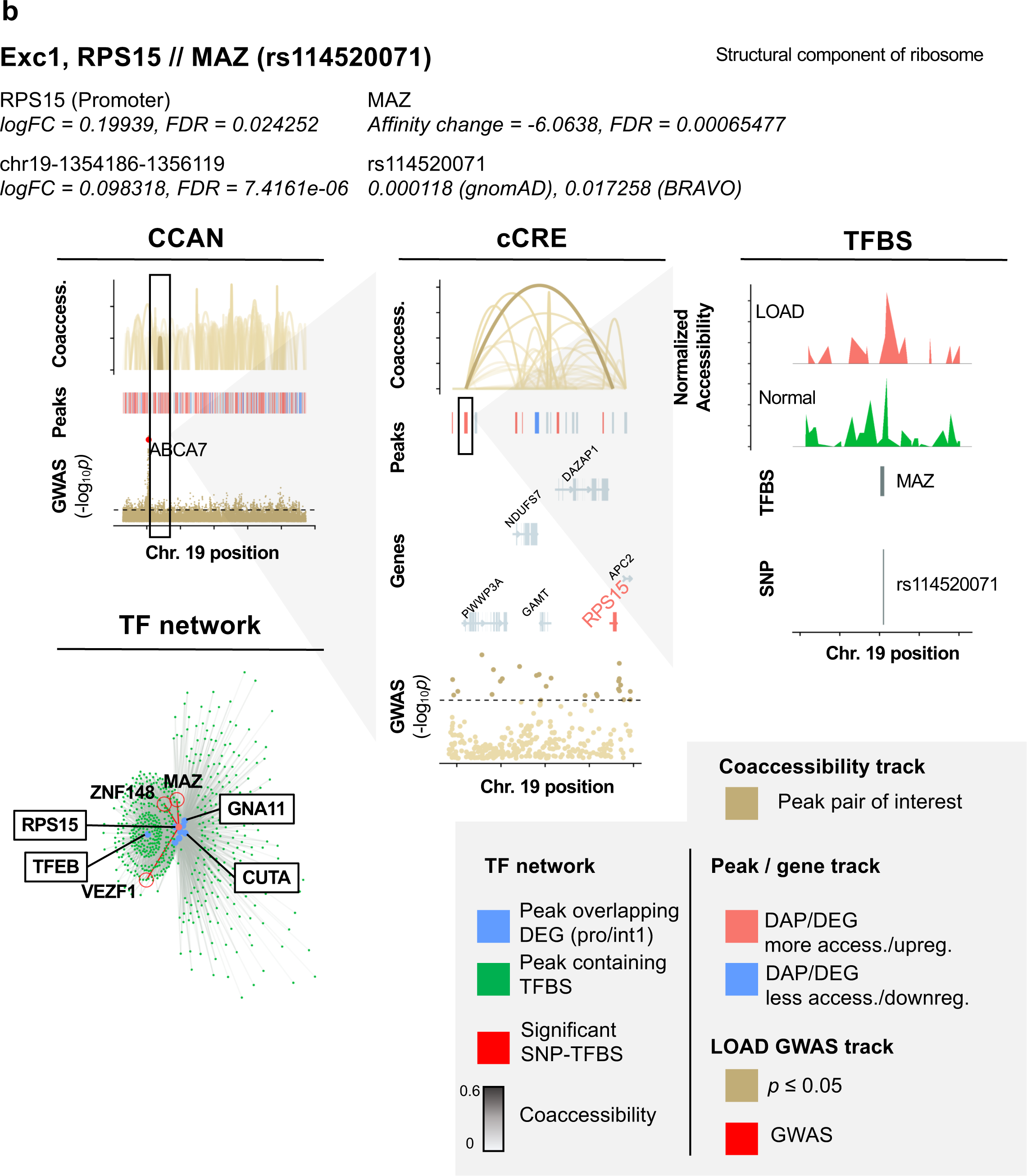
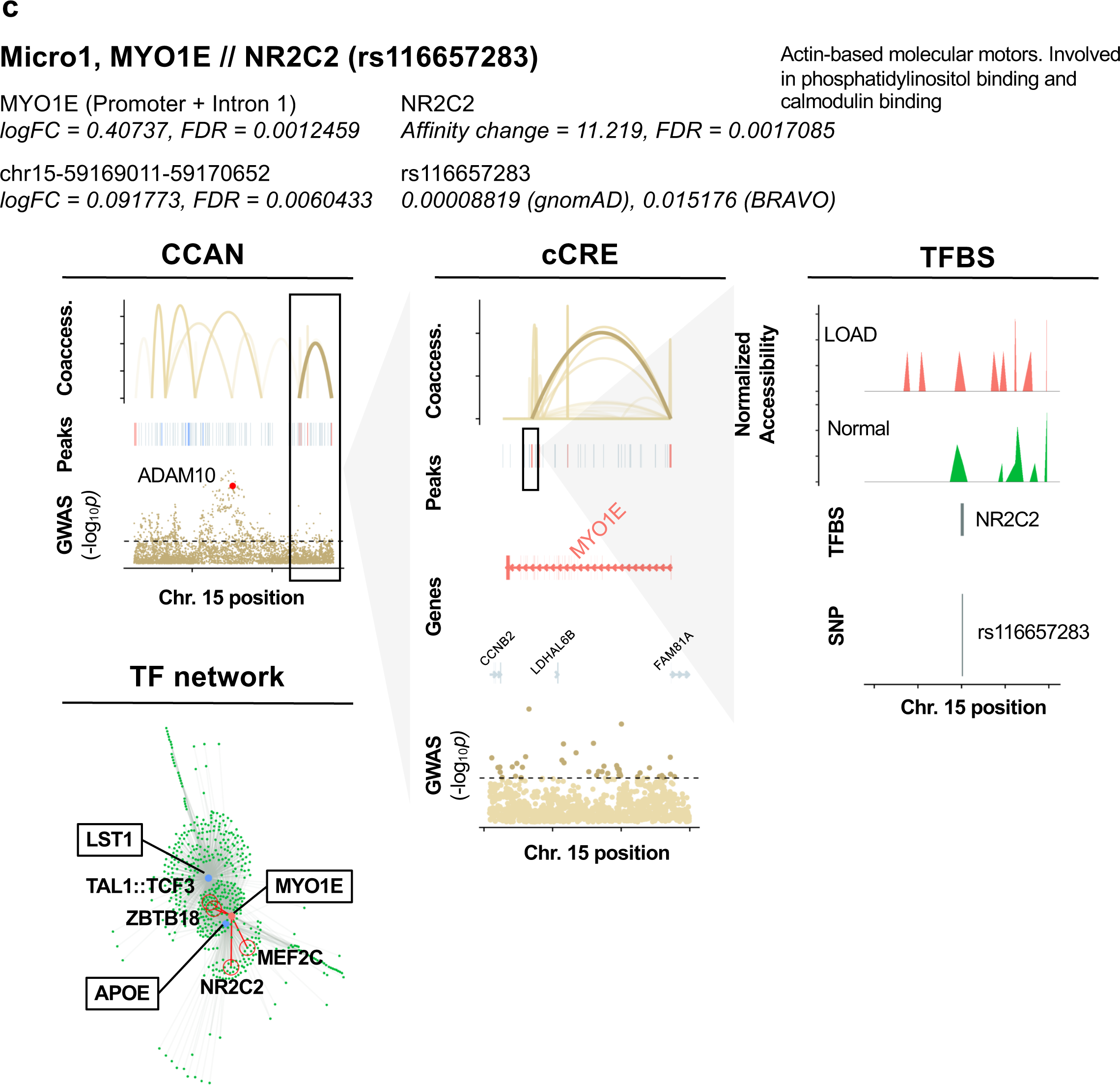
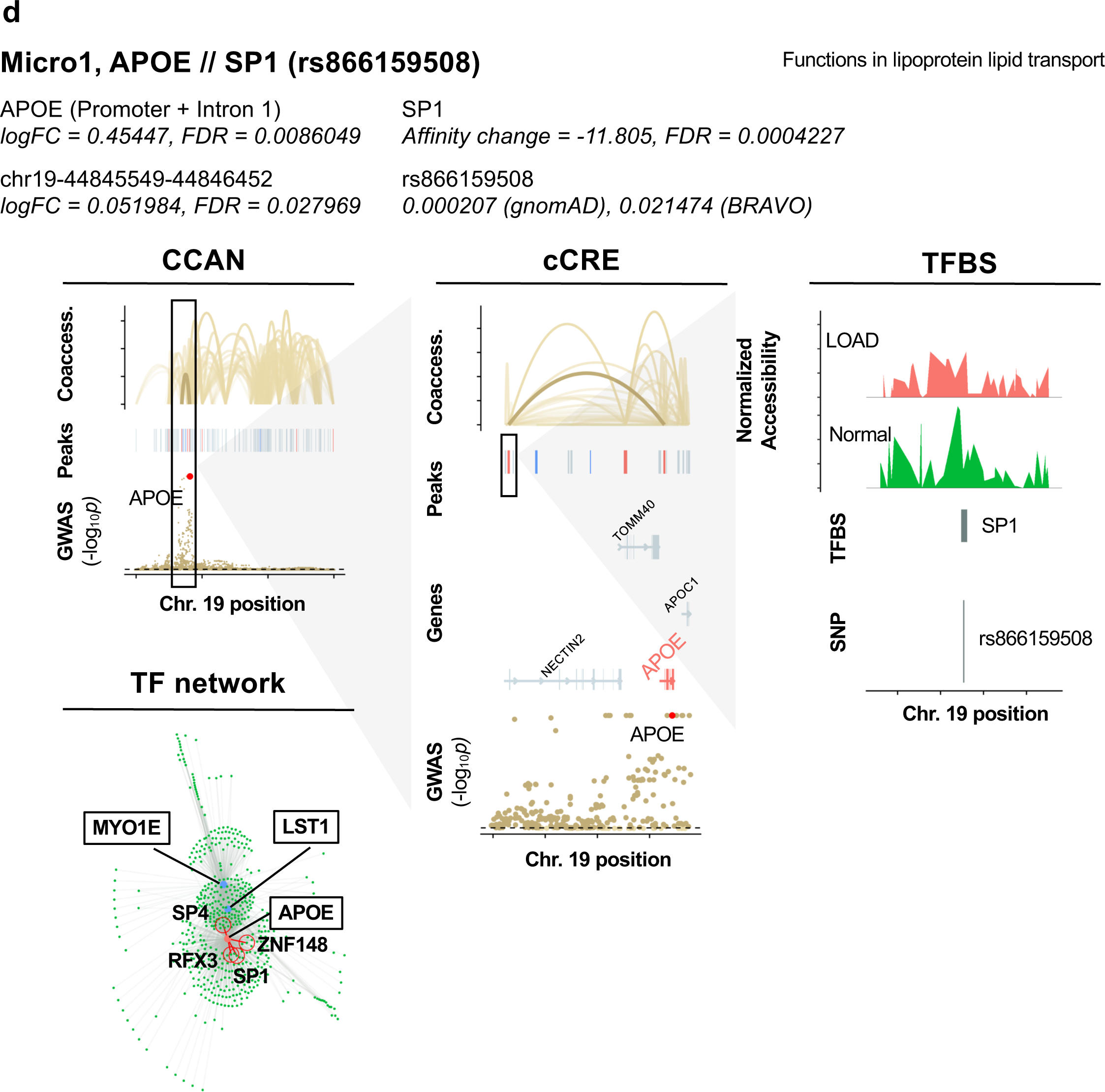
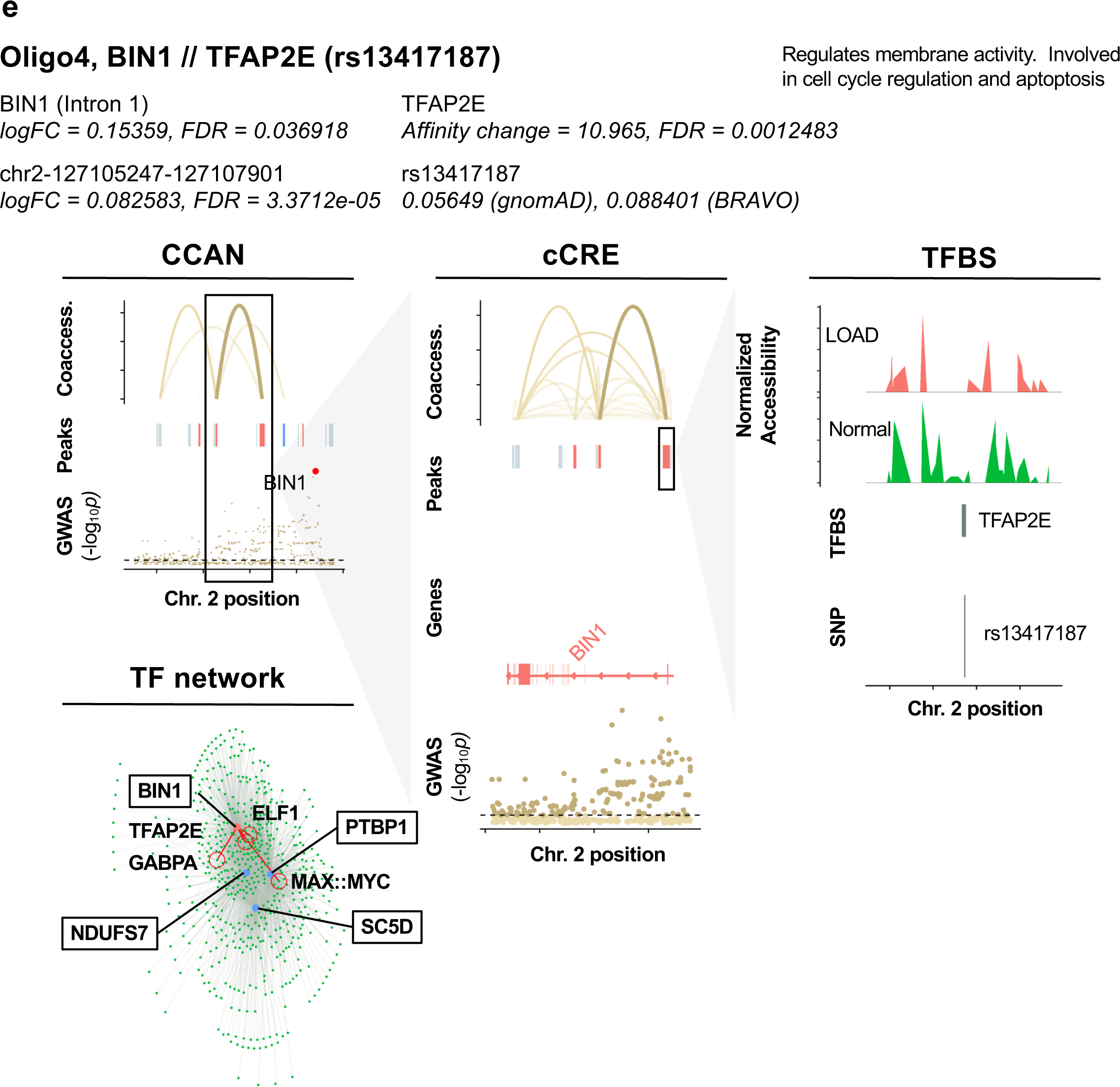
Identification of SNPs predicted to influence TF binding affinity at GWAS loci in LOAD CCANs. **a,** Summary tables of SNP-TFBS overlaps in unidirectional and mixed LOAD CCANs. **b-e**, Diagrams of specific example SNP-TFBS overlaps. The cell subtype, regulated DEG, TF and SNP ID are shown in bold. The log fold change (*Log2FC*) and probability values (*p*) are shown for each DEG and corresponding cCRE. Additionally, functional information for each DEG is provided. CCAN stacked plots show peak coaccessibility scores, directionality of changes in DAP accessibility in LOAD (red = increased accessibility, blue = reduced accessibility), and degree of LOAD association for GWAS loci. All features are arranged along the same horizontal access to indicate chromosomal position. cCRE stacked plots are detailed from boxed area of CCAN plots additionally indicate overlapped gene coding regions, with upregulated DEGs shown in red and downregulated DEGs shown in blue. TFBS activity stacked plots are detailed from boxed areas of cCRE plots and indicate normalized accessibility of genomic region in LOAD and normal samples as well as aligned chromosomal positions of TFBSs and SNPs. TF Network plots illustrate potential regulatory networks between DEG-overlapping peaks (blue) and TFBS-overlapping peaks (green), with those linkages predicted to be affected by LOAD SNPs shown in red.

## DISCUSSION

In this study, we performed *parallel* snATAC-seq and snRNA-seq using brain samples from European ancestry subjects and identified LOAD-associated transcriptome and chromatin accessibility signatures and their crosstalk at a cell subtype-level resolution. To our knowledge, this is the first study that identified candidate noncoding regulatory variants and their interacting TFs in LOAD by integration of single-nuclei multi-omics datasets, and provided evidence suggesting that LOAD genetic variants exert their putative pathogenic effects in a brain cell subtype-specific manner.

Previously, we applied ATAC-seq on NeuN sorted nuclei and identified multiple LOAD specific neuronal (NeuN+) and non-neuronal (NeuN-) chromatin accessibility sites, several of which overlapped with LOAD-GWAS regions. Furthermore, we identified sex-dependent LOAD changes in chromatin accessibility, particularly, glia-specific sites were found only in females. Last, by integrative analysis of the LOAD-GWAS regions, ATAC-seq on sorted nuclei, and snRNA-seq, we functionally characterized the impact of these chromatin accessibility differences on gene expression within LOAD loci ^66^. The current study extends our previous work in several ways. We now provide granular insight into cell subtypes molecular changes. Furthermore, we uncovered the crosstalk between epigenetic, genomic, and transcriptomic determinates of LOAD pathogenesis. Our outcomes provide catalogues of candidate genes, cCREs, and variants involved in LOAD genetic etiology and the cell subtypes in which they act to exert their pathogenic effects.

To date, only a few other studies have profiled gene expression in cortex tissue of LOAD patients at single-cell resolution ^34–37, 67–69^, out of which only two studies have characterized both the transcriptome and chromatin accessibility of LOAD ^36, 37^. These pioneering studies of snRNA-seq from cortexes of LOAD patients found that the strongest LOAD-associated changes appeared early in pathological progression and were highly cell-type specific ^34^, and identified LOAD-associated gene dysregulation in specific cell subpopulations, particularly for *APOE* and transcription factor genes ^35^. One study used these datasets to examine the sex dependent effects of LOAD and found disproportionate representation in disease-associated cell subtype populations as well as differential LOAD expression profiles between the sexes ^34^, while another characterized markers of selectively vulnerable neuron populations in AD ^68^. Nonetheless, these studies focused only on snRNA-seq data. More recently, an integrative multi-omics framework of snRNA-seq and snATAC-seq in late-stage LOAD nuclei identified disease-associated cCREs and analyzed gene coexpression networks ^36^. Another study examined gene expression and chromatin accessibility within the same nuclei and identified candidate transcription factors regulating LOAD-associated gene expression in neurons and microglia ^37^. Altogether, our and others’ studies have demonstrated the importance of cell-type and subtype specific omics profiling of human brain tissues to advance the understanding of the molecular and cellular subtype specific pathways underlying LOAD. Data sharing across groups will not only provide accessibility to replication cohorts but will also further the genetic exploration using meta-analysis in larger sample size to validate discoveries and test different hypotheses.

While GWAS infer genes based on proximity to the most strongly associated SNP, identification of the actual genes and variants involved in disease risk has been a challenge. We examined 1 Mb regions surrounding LOAD tagging SNPs and provided multiple examples for DEGs, *i.e.* candidate genes involved in LOAD, that mapped within the LOAD associated region but more distal from the tag SNP. Additionally, we performed the first step towards identification of candidate regulatory SNPs and their linked genes. Using single-nucleus genomics integrative analysis focused on LOAD GWAS loci, we prioritized several SNPs for validation studies. Notably, the associations of these candidate SNPs with LOAD risk remain to be determined. In future studies, a larger sample size may allow conducting of transcriptomic and chromatin accessibility QTL/eQTL mapping to determine colocalization with GWAS loci.

The importance of microglia in AD has been previously established ^70–72^. Cells of this type may have a protective function in β-amyloid (Aβ) clearance via phagocytosis and protease degradation in the early stages of the disease ^73, 74^, but may also contribute to Aβ plaque accumulation in later stages via seeding and proinflammatory cytokine production ^75–78^. Microglia have also been shown to exacerbate neuronal synapse loss, neurotoxic inflammation, and tau pathology in AD ^79–82^. Furthermore, a high number of identified AD risk genes have been found to be preferentially expressed in microglia ^83^. Notably, 22% of AD GWAS loci-proximal genes identified by Bellenguez et al. ^8^ were also microglial signature genes. It was further shown via ChIP-seq and ATAC-seq that AD-associated non-coding variants are specifically enriched in microglial enhancer regions over those of other brain cell types ^12^. Both *APOE* and *MYO1E* expression have been previously associated with disease-associated microglia exhibiting neurodegeneration-specific gene expression profiles in mouse models of AD ^56^. Expression of human *APOE*e4 also resulted in increased tau pathology and inflammatory microglial response in mice, and increased production of TNFα by microglia *in vitro* ^84^. *APOE* has been further implicated in conjunction with the microglia-associated receptor protein *TREM2* in driving the conversion of microglia to a neurodegenerative phenotype ^85^. In this regard we would like to highlight our findings with *APOE* and *MYO1E*. We identified both *APOE* and *MYO1E* as key LOAD-associated genetic factors within microglia in several distinct analyses. First, these genes were the most strongly LOAD-upregulated GWAS DEGs in multiple microglial subtypes (Fig. 3c) as well as microglia overall (Fig. 3e), while genes associated with mediation of synaptic processes were strongly downregulated among microglia generally (Fig. 3f). Furthermore, *APOE* and *MYO1E* were also both linked to cCREs within LOAD CCANs of the Micro1 cluster (Fig. 5b, d), and moreover atSNP analysis revealed multiple SNPs predicted to impact TF binding affinity within cCREs potentially regulating both of these genes within this same cell subtype. Taken together, these findings serve to further underscore the importance of *APOE* and *MYO1E* as LOAD risk factors that exert their pathogenesis in microglia, and also suggest potential molecular mechanisms for dysregulation in LOAD.

Over the last decade, LOAD GWAS have confirmed strong associations with the *APOE* LD genomic region, and no other LOAD-association remotely approached the same level of significance ^1, 3, 7, 86–90^. However, whether the strongest signal is attributed to additional variants and haplotypes within this LD region jointly with e4, as well as the molecular mechanisms underlying the LOAD-association with the APOE LD region, is largely unknown. While *APOE*e4 is the first and strongest genetic risk for LOAD, accumulating evidence has suggested that the increased overall expression of *APOE* plays an important role in the etiology of LOAD (reviewed by Gottschalk et al. ^91^ and Yang et al. ^92^). Foremost, previously we found significantly higher levels of *APOE*-mRNA in brain tissues obtained from e3/3 LOAD patients compared to 3/3 healthy donors, consistently with other reports showing elevated levels of *APOE*-mRNA in LOAD brains ^14–16, 93^. In addition, new snRNA-seq datasets showed LOAD changes in *APOE* expression in glial cell-types, in particular upregulation in microglial subpopulations ^34, 35, 94^. Moreover, studies using the *APP/PS1* transgenic mice showed that lowering the ApoE protein levels ameliorated cognitive dysfunctions and Aβ pathology ^95^ independent of the *APOE* allele ^96–98^. Lastly, studies showed LOAD associated differential DNA-methylation ^29, 99–102^, further supporting a role for dysregulation of *APOE* expression in the genetic etiology of LOAD. Here, to circumvent the confounding effect of *APOE*e4, we constrained the analysis to *APOE*e3 only. We observed both increased expression and increased chromatin accessibility of *APOE* among microglial populations. Additionally, we identified multiple cCREs linked to both the promoter and intron 1 sequences of *APOE* in the Micro1 subtype cluster, as well as multiple SNPs potentially influencing TF binding to the *APOE* promoter in this same cluster, suggesting a possible mechanism for microglial *APOE* dysregulation in LOAD. Our results thus provide further evidence that there are clear changes in *APOE* expression associated with LOAD and independent of the e4 allele^96^, suggesting that regulation of *APOE* expression in specific cell subtypes may impact the risk to develop LOAD, making the modulation of the overall ApoE protein levels useful as a future therapeutic target.

Despite major advances in genome technology and our innovative experimental approach and analytical strategy, there are still a number of limitations associated with the analysis of single cell data. In this study, for cell type annotation and subtype clustering, variance stabilizing transformation (SCT) ^103^ of the snRNA-seq count data was employed. While SCT offers improvement over log normalization by regularizing the effect of sequencing depth on transcript counts, it results in data too computationally intensive to use for differential expression analysis. Likewise, the term frequency-inverse document frequency (TF-IDF) normalization ^104^ used in snATAC-seq clustering addresses some of the pitfalls of log-normalization, but in some cases these corrections may be too harsh. A regularized regression framework may offer more reproducible results, but again would require prohibitively intensive processing power. These methods should be possible in future work with greater computational resources. Another limitation involves the accuracy of cluster linking. For each matching snATAC-seq and snRNA-seq cluster, we found that the results showed an overall 75% accuracy rate. The most accurate cluster in the test dataset had a Jaccard index of 0.95, while the least accurate cluster had a Jaccard index of 0.32. Despite this limitation, in all clusters, the cell type of the closest matching snRNA cluster was concordant with the initially predicted cell type of the snATAC cluster. Moreso, the intrinsic structure of the snATAC clusters was not lost. Thus, the downstream analysis remained unaffected while providing loosely informative insight towards epigenetic and transcriptomic interplay. Finally, previous research has estimated that a high proportion of predicted TF binding sites are false positives^105–107^. Raising the stringency of *p*-value thresholds decreases the rate of false positive motif matches, but increases the rate of false negative matches even more strongly, thus limiting the value of this approach ^108^. We attempted to address this limitation by restricting the analysis to TFs that were expressed in ≥10% of the corresponding cell subtype, so that our findings presented a biological relevance. Currently, single cell methods such as ChIP-seq or ChIA-PET-seq^109^ are not available to validate our findings experimentally. Similarly, chromatin confirmation capture (3C) or chromatin confirmation capture carbon copy (5C) datasets on temporal cortex or analogous tissue regions in a single cell-type resolution would provide the most direct evidence for the cCRE-DEG interactions identified in our study, as they become available.

Single cell analyses specifically for disease-involved heterogenous organs such as the brain are imperative. Molecular changes are cell subtype specific, meaning that a particular gene and/or variant may exert its effect on a certain cell subtype but can be neutral in another cell subtype. LOAD exemplifies a genetically complex disease affecting a heterogenous tissue and therefore requires granularity in research approaches to facilitate advancements and new discoveries in the field. Our study has pioneered the powerful strategy of integration of cell-type specific multi omics datasets collected from the same sample in parallel to describe *cis-trans* regulatory networks disrupted in LOAD, validate known LOAD loci, and identify new candidate genes. We have furthermore presented the most comprehensive interrogation to date of genetic variants potentially impacting gene regulatory networks in LOAD. Further investigations including sex and ancestry stratified studies using integrative single cell multi omics data will advance our understanding of the genetics underpinning LOAD in specific populations. Collectively, our findings provide a rich dataset for future mechanistic experiments, confirm known LOAD GWAS loci while also identifying novel loci, and highlight the disease-relevant cell types and subtypes for follow-up validation studies in model disease systems and for development of future therapeutic interventions.

## CONCLUSIONS

Profiling the chromatin accessibility and transcriptomic landscapes from the same pool of nuclei and at the same time is a well-controlled approach to facilitate multi-omic integrative analyses and advance new genetic discoveries in complex disorders such as LOAD. Our study has seven major findings for the field of LOAD genetics. First, we have generated cell subtype-specific profiles of LOAD-associated chromatin accessibility signatures and maps of LOAD cCRE. Second, we leveraged the LOAD accessible peaks dataset to identify candidate co-accessible networks in each cell subtype. Third, we provided a catalogue of candidate LOAD-associated cell subtype specific DEGs. Fourth, we identified TFs relevant to LOAD and the cell subtype in which they act. Fifth, we catalogued candidate SNPs involved in dysregulation of key genes in LOAD in a cell subtype specific manner. Sixth, we have demonstrated that LOAD associations may not be interpreted by the most proximate gene. Seventh, we provided evidence that the genetics underpinning LOAD risk mediates its pathogenic effects in various glial and neuronal cell subtypes. Overall, these results suggest that cell subtype-specific *cis-trans* interactions between regulatory elements, noncoding variants and TFs, and the genes dysregulated by these networks contribute, at least in part, to the development of LOAD.

## METHODS

### Human post-mortem brain tissue samples

Frozen human temporal cortex tissue of LOAD samples (*n* = 12) and neurologically healthy control samples (Normal) (*n* = 12) was obtained from the Kathleen Price Bryan Brain Bank (KPBBB) at Duke University. The demographics for this cohort are included in Table 1 and detailed in Table S1. Clinical diagnosis of LOAD was pathologically confirmed using Braak staging (AT8 immunostaining) and amyloid deposition assessment (4G8 immunostaining) for all LOAD samples. All tissue donors were Caucasians with the *APOE* e3/e3 genotype and Braak & Braak Stage III-V. The project was approved for exemption by the Duke University Health System Institutional Review Board. The methods described were conducted in accordance with the relevant guidelines and regulations.

### Cohort Statistics

For comparisons of demographic variables, R statistical programming language was used. Age and post-mortem interval (PMI) of female LOAD was compared to female Normal, and age and PMI of male LOAD was compared to male Normal. The Shapiro-Wilk test was used for normality, Bartlett’s test for equal variance of normally distributed data, and Levene’s test for equal variance of non-normally distributed data. If groups were normal and had equal variance, two sample *t*-tests assuming equal variances were used to determine differences between group means. If groups were not normal, a Mann-Whitney’s *U* test was run. No groups had unequal variances.

### Nuclei isolation from post-mortem human brain tissue

The nuclei isolation procedure was based on previous studies^110, 111^, but has been optimized for single-cell experiments. 100-200 mg of post-mortem human brain tissue samples (gray matter) was thawed over ice in Lysis Buffer (0.32 M Sucrose, 5 mM CaCl_2_, 3 mM Magnesium Acetate, 0.1 mM EDTA, 10 mM Tris-HCl pH 8, 1 mM DTT, 0.1% Triton X-100) and homogenized using a 7 ml dounce tissue homogenizer (Corning) with pestle A. The homogenate was filtered through a 100 μm cell strainer, transferred to a 14 x 89 mm polypropylene ultracentrifuge tube, and carefully underlain with sucrose solution (1.8 M Sucrose, 3 mM Magnesium Acetate, 1 mM DTT, 10 mM Tris-HCl, pH 8). The nuclei were separated by ultracentrifugation at 4°C at 107,000 RCF for 15 minutes. The supernatant was removed by aspiration, and the remaining nuclei were washed with 1 ml Nuclei Wash Buffer (10 mM Tris-HCl pH 8, 10 mM NaCl, 3 mM MgCl_2_, 0.1% Tween-20, 1% BSA, 0.2 U/μL RNase Inhibitor) and incubated on ice for 5 minutes. The nuclei were gently resuspended, and 800 μL was transferred to a microcentrifuge tube designated for the 10X Genomics single-cell ATAC assay while 200 μL was transferred to a microcentrifuge tube designated for the 10X Genomics single-cell gene expression assay. The nuclei were centrifuged at 300 RCF for 5 minutes at 4°C, and the supernatant was again aspirated. For the ATAC assay, the pellet was resuspended in Diluted Nuclei Buffer (10X Genomics). For the gene expression assay, the pellet was resuspended in Wash and Resuspension Buffer (1X PBS, 1% BSA, 0.2 U/μL RNase Inhibitor). After a 1-minute incubation on ice, the nuclei were filtered through a 35 μm strainer. Nuclei concentrations were determined using a Countess™ II Automated Cell Counter (ThermoFisher) and nuclei quality was assessed at 10X and 40X magnification using an Evos XL Core Cell Imager (ThermoFisher) prior to library construction.

### Parallel snATAC-seq/snRNA-seq library preparation and sequencing

Single-nucleus (sn)ATAC-seq libraries were constructed using the Chromium Next GEM Single Cell ATAC Library and Gel Bead v1.1 kit, Chip H Single Cell kit, and Single Index Kit N Set A (10X Genomics) according to manufacturer’s instructions. In parallel, from the same pool of nuclei from each sample, single-nucleus (sn)RNA-seq libraries were constructed using the Chromium Next GEM Single Cell 3’ GEM, Library, and Gel Bead v3.1 kit, Chip G Single Cell kit, and i7 Multiplex kit (10X Genomics) according to manufacturer’s instructions. For each sample, 10,000 nuclei were targeted for both the ATAC and 3’ assays. Library quality control was performed on a Bioanalyzer (Agilent) with the High Sensitivity DNA Kit (Agilent) according to manufacturer’s instructions and the 10X Genomics protocols. Libraries were submitted to the Sequencing and Genomic Technologies Shared Resource at Duke University for quantification using the KAPA Library Quantification Kit for Illumina® Platforms and sequencing. Groups of four snATAC-seq libraries from 1 LOAD female, 1 LOAD male, 1 Normal female, and 1 Normal male were pooled on a NovaSeq 6000 S1 100bp PE full flow cell to target a sequencing depth of 400 million reads per sample (Read 1N = 50, i7 index = 8, i5 index = 16, and Read 2N = 50 cycles). Groups of four snRNA-seq libraries from 1 LOAD female, 1 LOAD male, 1 Normal female, and 1 Normal male were pooled on a NovaSeq 6000 S1 50bp PE full flow cell to target a sequencing depth of 400 million reads per sample (Read 1 = 28, i7 index = 8, and Read 2 = 91 cycles). Sequencing was performed blinded to diagnosis, age, and sex.

### snRNA-seq data processing

Raw sequencing data from snRNA-seq experiments was converted to fastq format, aligned to a GRCh38 pre-mRNA reference, filtered, and counted using CellRanger 4.0.0 (10X Genomics). Subsequent processing was done using Seurat 4.0.1^112^. Filtered feature-barcode matrices were used to generate Seurat objects for the 24 samples. For quality control filtering, nuclei with less than 200 or greater than 10,000 features were excluded. Nuclei with greater than 17.4% mitochondrial gene expression were found to cluster together on a uniform manifold approximation and projection (UMAP) feature plot and were also excluded. Because experiments were conducted in nuclei rather than cells, mitochondrial genes were subsequently removed. The 24 Seurat objects were merged into one, and were iteratively normalized using SCTransform^113^ with glmGamPoi, which alleviates bias from lowly expressed genes^114^. Batch correction was performed using reference-based integration^38^ on the 24 normalized datasets, which improves computational efficiency for integration.

Cell type annotation was conducted using a label transfer method^38^ and a previously annotated reference dataset from human M1 (see below). Batch corrected data from both our dataset and the human M1 dataset were used for label transfer. Nuclei with maximum prediction scores of less than 0.5 were filtered out. Nuclei with a percent difference of less than 20% between their first and second highest cell type prediction scores were termed “hybrid” and subsequently removed^35^. Endothelial cells and vascular leptomeningeal cells (VLMCs) were in low abundance (465 total) and did not form distinct clusters in UMAP analysis and were therefore filtered out of the final dataset. After running a Principal Component Analysis (PCA), dimensionality was examined using an Elbow plot and by calculating the variance explained by each principal component (PC). UMAP analysis was then run with the first 30 PCs, and nuclei were clustered based on the UMAP reduction at a resolution of 0.1. Counts of predicted cell types based on the label transfer were examined for each of the 33 clusters (Table S4), and clusters were manually annotated based on the majority cell type for each cluster (e.g., ‘Exc1’, ‘Exc2’, etc.).

### Human M1 reference data processing

A recent snRNA-seq study on human primary motor cortex (M1) used 10X Genomics technology to characterize 127 transcriptomic cell types^33^. To optimize label transfer, we re-processed the data to map it to GRCh38 Ensembl 80 as we did with our data. Fastq files were obtained from the Neuroscience Multi-omic Data Archive (NeMO: https://nemoarchive.org/) and were aligned to the same GRCh38 pre-mRNA reference used for our data, filtered, and counted using CellRanger 4.0.0 (10X Genomics). Filtered feature-barcode matrices were used to generate 14 Seurat objects, and nuclei that were absent from the annotated metadata from the study were filtered out, leaving 76,519 nuclei in the final re-processed dataset. The Seurat objects were merged and iteratively normalized using SCTransform^113^ with glmGamPoi. Batch correction was performed using reference-based integration^38^ on the 14 normalized datasets. The 127 transcriptomic cell types were grouped into 8 broad cell types including astrocytes, endothelial cells, excitatory neurons, inhibitory neurons, microglia, oligodendrocytes, oligodendrocyte precursor cells (OPCs), and VLMCs.

### Cell type proportion comparisons

To compare proportions of cell types and subtypes between LOAD and control groups, nuclei of each type and subtype were counted and divided by the total nuclei for each sample. Then, a bootstrapped two-sided Wilcoxon rank-sum test was performed in R (v4.0.2) using the wilcox.test function with default parameters and Benjamini–Hochberg correction for multiple testing. 20% of nuclei were randomly selected from all samples under comparison in each of 30 iterations^36^.

### snATAC-seq data processing

DNA fragments acquired from our snATAC-seq experiments were sequenced and converted to fastq format, from which they were mapped to GENCODE’s human release 32 reference^115^ and counted using CellRanger-ATAC 1.2.0 (10X Genomics). We screened the remaining nuclei using the following quality control metrics:

a. Nucleosome signal: Defined as the ratio of mononucleosome fragments (147 to 294 bp) to nucleosome free fragments (< 147 bp). Nuclei having a nucleosome signal of greater than 4 were removed^116^.
b. Transcription start site (TSS) enrichment: The ratio of aggregated, normalized read signal centered around a reference set of TSS’s compared to the signal in the TSS flanking regions. Nuclei with a TSS enrichment score of less than 2 were removed^116^.
c. Percent reads in peaks: The proportion of fragments in the cell that map to peak regions. Cells with less than 15% of reads in peaks were removed^116^.
d. Total peak region fragments: Cells with less than 1000 peak region fragments were discarded due to low sequencing depth. Additionally, cells in the upper 1% in each sample distribution were removed as a precaution against multiplets^116^.
e. Blacklist ratio: The proportion of fragments that map to sequences associated with technical artifacts. Cells with greater than 5% of fragments mapping to blacklisted regions were removed^117^.

The remaining preprocessing steps were conducted using R packages Seurat 4.0.1^38^, Signac 1.3.0^116^, and Harmony 0.1.0^118^. Latent semantic indexing (LSI) was used to create a low-rank approximation of the data^119^. The 24 datasets were term frequency-inverse document frequency (TF-IDF) normalized and aggregated to form a joint peak by cell count matrix. Then, singular value decomposition (SVD) was performed on the joint dataset, after which the left singular vectors were standardized, representing the LSI components. Then, the correlation of each component with sequencing depth was measured. Due to high correlation with sequencing depth, the first dimension was removed from downstream analysis (rho = 0.7)^116^. The remaining LSI components were then adjusted with the function RunHarmony to remove batch effects before clustering the data. In alignment with snRNA dimensionality reduction methods, we used dimensions 2 through 30 for clustering and downstream analysis.

Clusters were constructed from the adjusted LSI embeddings of the integrated dataset using the Seurat functions FindNeighbors and FindClusters, with *k*-nearest neighbors set to 20, and a cluster resolution of 1. The data was then projected onto a 2D surface using the uniform manifold approximation (UMAP) algorithm and inspected to ensure ample cluster resolution.

### Cell type annotation of snATAC-seq nuclei

Cell type annotation of the snATAC cells was done using the integrated snRNA dataset as a reference^38^. First, “gene activity” matrices were constructed from each snATAC sample by counting the fragments mapping to promoter regions (between 2000 bp upstream and 200 bp downstream of TSS) of each cell. After quantifying promoter region fragments, the matrices were log-normalized using Seurat.

The Seurat function FindTransferAnchors was used to annotate the snATAC data against our snRNA data, utilizing canonical correlation analysis as the reduction method. Seurat computes the cross correlation between variable features of snATAC and snRNA cells. After L2 normalization, the left and right singular vectors from the SVD of this matrix are taken as the canonical correlation vectors. Seurat then uses a mutual nearest neighbor approach to find anchors between the datasets, representing biologically similar cell states across modalities. For each cell, the weighted combination of the *k*-nearest anchors was used to calculate prediction scores for each of the major cell types. For each cell, the predicted cell identity was the cell type with the maximum prediction score. Nuclei with maximum prediction scores of less than 0.5 were filtered out. “Hybrid” nuclei were identified using the same metric as for snRNA-seq data above, and subsequently removed.

### Linking snATAC and snRNA datasets

The Seurat functions *FindTransferAnchors* and *TransferData* were used in a comparable manner to link snATAC clusters to snRNA clusters. As in cell type annotation, the anchors were used to transfer the snRNA cluster information to the snATAC cells. Each cell in the snATAC data was given 33 prediction scores, corresponding to each of the snRNA clusters. Directly clustering snATAC cells by using the snRNA cluster prediction scores did not fully align with the intrinsic structure of the snATAC data found from the initial Louvain clustering. As such, the original snATAC clusters were left unchanged, and the snRNA cluster prediction scores in each cell were summed across all nuclei belonging to the same snATAC cluster. The maximum prediction score in each snATAC cluster was used to designate a closest matching snRNA cluster. Lastly, the cell type of the linked snRNA cluster was matched against the original cell type of the snATAC cluster to ensure concordance.

To assess the accuracy of cluster linking, we used PBMC granulocyte multiome data, freely available on the 10X Genomics website. The snATAC and snRNA assays were processed separately using the same pipelines outlined in this manuscript (quality control, normalization, and community detection). Each snATAC cluster was designated a closest matching snRNA cluster by summing prediction scores of each cell as described previously. For each matching snATAC and snRNA cluster pair, the Jaccard index of the cluster barcode compositions was calculated. Additionally, the hybrid score for each snATAC cluster was calculated as described in cell type annotation, using the overall cluster prediction scores as input. Cluster hybrid scores provided a measure of the consensus amongst cells in each snATAC cluster.

### Peak Calling

As different cell types exhibit epigenetic heterogeneity, it was necessary to perform peak calling on each cluster before differential accessibility analysis to reduce noise and computational burden. Peak regions were predicted empirically using the MACS2 algorithm on each cluster within each sample separately^120^. MACS2 assumes a dynamic Poisson distribution, with λ dependent on the read signal of the surrounding genomic region and is robust to technical and biological bias. Peaks were designated as regions having a *p* value of ≤ 10e-5. To combine peaks into a consensus set for each cluster, Multi Sample Peak Calling (MSPC) software package was used^121^, which employs Fisher’s method to evaluate overlapping peaks across samples. Peaks that occurred in at least 2 samples, with an FDR of ≤ 0.05 from Fisher’s combined probability test were used as the consensus set for downstream analysis.

### Covariate selection for differential analyses

Prior to differential analysis, we assessed the potential impact of several technical variables from each of the snRNA-seq and snATAC-seq experiments separately such as number of nuclei, sequencing saturation, and reads mapped to the genome, as well as donor-level characteristics such as age, sex, and PMI. Several processing steps were taken prior to association testing on the snRNA-seq and snATAC-seq data separately. For each experiment, read counts were summed for all nuclei per donor sample, resulting in only one expression or accessibility peak value per donor sample per gene or chromatin peak, respectively. This down-coding was done for the covariate selection analysis to address the fact that all nuclei from each donor would have identical donor characteristics. Subsequently, genes with no expression or peaks with zero accessibility for >20% of samples were removed, and all values were mean centered and scaled prior to analysis.

Principal component (PC) analysis was performed using prcomp in R for all genes and peaks passing our pre-processing steps, separately. We then performed linear regression using glm in R of PCs explaining >10% of the variability in global expression or chromatin accessibility on both nuclei- and donor-specific metadata variables to identify factors that should be included as covariates in differential analysis. Specifically, we selected the variable most associated (surpassing Bonferroni correction for multiple testing, *q*<0.05) with PC1 (or alternatively, the PC explaining the most variability) and regressed all genes or peaks on the associated variable to obtain gene or peak residuals that are adjusted for its effect. We then performed PC analysis on the gene or peak residuals, and in an iterative process, repeating the above steps until no additional metadata variables were associated with global expression or chromatin accessibility (*q*<0.05). For the snRNA-seq analysis, sex, age, PMI, sequencing saturation, and cluster proportions (calculated for each donor by dividing the number of nuclei of each cell type or subtype by the donor’s total nuclei count) were selected as covariates for the differential analysis, and for the snATAC-seq analysis, sex, age, PMI, peak region fragments, cluster proportions, and percent fragments overlapping any targeted region were selected.

### Differential expression analysis

To avoid pseudoreplication bias, we used MAST^122^ with a random effect for donor, as in a recent publication^43^. For each cell type and cluster, raw counts from the snRNA-seq assay were log_2_(*x* + 1) transformed, and genes expressed in less than 10% of cells in either group (LOAD or Normal) were filtered out. For differential expression testing, a two-part hurdle model was run, consisting of a zero-inflated regression fitting a generalized linear mixed-effects model followed by a likelihood ratio test comparing the model with and without the group factor. The reference level was set to ‘Normal’ such that the results for logFC coefficients would be positive if up-regulated in LOAD and negative if down-regulated in LOAD. Cellular detection rate (number of genes expressed) was calculated for each nucleus, centered, and scaled, and added to the model as a covariate to control for nucleus size. The proportion of nuclei for each cell type and cluster was calculated for each donor (e.g., for a given cell type, the number of nuclei for a donor divided by that donor’s total nuclei count) and added to the model to control for sample-to-sample variation in cell type composition. *P* values were adjusted for false discovery rate (FDR) to correct for multiple comparisons. The percentage of nuclei expressing each gene was calculated for both groups and added to the results.

### Differential accessibility analysis

Seurat’s “LR” test was used for differential accessibility testing between LOAD and normal samples was performed on each cell type and each cluster ^116, 123^. The model predicted diagnosis using a binomial regression, with peak region fragments, percent fragments overlapping any targeted region, nuclei proportion, age, sex, PMI, and fragment counts as the predictors. A likelihood ratio test was conducted to compare the null and experimental models, using fragment counts as the parameter of interest. Corresponding *p* values for each peak were adjusted for FDR on a per cluster basis. As with snRNA-seq data, positive log fold change corresponded to increased accessibility, and vice versa.

### Assessment of *cis* co-accessibility networks

Next, we sought to characterize chromatin interactions in the data using the R package Cicero^44^. The Cicero pipeline was conducted on a per cluster basis using LOAD cells only. We passed our integrated LSI embeddings to Cicero’s bootstrap aggregation procedure, in which highly similar cells are aggregated by summing the raw counts in groups of 50 *k*-nearest neighbors. The fragment sums are then normalized to account for within-group sequencing depth. Cicero then uses a graphical LASSO to estimate the partial correlation structure of each peak with its neighboring peaks. A penalty term dependent on the genomic distance between peak pairs is used in GLASSO, and the resulting regularized correlations derived from the precision matrix are termed “co-accessibility scores.” We defined the maximum peak-peak distance, at which regularized correlations are assigned 0, as 500 Kbp. We then specified a minimum co-accessibility score of 0.2 before extracting *cis* co-accessibility networks (CCANs) from the resulting data using Louvain community detection.

### LOAD cCRE’s and GO analysis

To probe candidate cis-regulatory elements within CCANs, we first isolated CCANs that (a) contained ≥ 1 DAP and (b) contained ≥ 1 peak that overlaps the promoter or intron1 of a DEG in the closest matching snRNA cluster. An additional factor to consider was whether the direction (i.e., the sign of DAP and DEG log fold change) aligned within DAP-DEG pairs. This assessment was done on each CCAN individually. First, all DAPs in the CCAN were extracted. Second, all DEGs in which the promoter or intron 1 was located within a peak belonging to the CCAN were extracted. Third, all pairs of DAPs and DEGs were screened as to whether they were perturbed in the same direction. Based on this screening, the CCANs were then given the following designations:

a. *Unidirectional*: Of all possible DAP-DEG pairs in the CCAN, all had the same sign for log fold change.
b. *Mixed*: Of all possible DAP-DEG pairs in the CCAN, there was ≥ 1 pair that had the same sign for log fold change, but not all pairs had the same sign for log fold change.
c. *Bidirectional*: of all possible DAP-DEG pairs in the CCAN, there were no pairs that had the same sign for log fold change.

As we were primarily interested in enhancer promoter interactions, only unidirectional and mixed CCANs were considered. Within unidirectional and mixed CCANs separately, we looked for DAP-DEG overlap peak pairs that met the following criteria:

a. DAP was highly coaccessible (coaccessibility score of ≥ 0.2) with a peak overlapping the promoter or intron 1 of a DEG.
b. The peak overlapping the promoter or intron 1 of a DEG was also a DAP.
c. Both DAPs and the overlapping DEG had the same direction of effect.
d. DEG log fold change ≥ 0.15.

For each cluster, we pooled the overlapping DEGs from both unidirectional and mixed CCANs that met these criteria and performed GO analyses with the R package topGO^124^. Genes expressed in at least 10% of cells in their corresponding cluster were used as the background gene sets. We used Fisher’s hypergeometric test statistic in evaluating enrichment of GO terms. Terms of interest were defined as having a *p* value ≤ 0.05 and mapping to at least 3 genes in the test gene set.

### Motif detection and enrichment analysis

To detect transcription factor binding sites (TFBS) in the data, position weight matrices (PWMs) from JASPAR 2020 were used to scan the genome for motifs within the CCANs of each cluster^125, 126^. The *p* value threshold for a motif match was 5e-5. For overlapping motif matches of the same transcription factor, only the highest scoring match was used. Given the high rate of type I error associated with *in silico* motif discovery algorithms, motif enrichment was also performed to rule out unlikely candidates. We used HOMER to detect motif enrichment in CCANs^127^. HOMER first quantifies the GC content and *n*-mer composition of both the background and target regions and applies weights to eliminate sequence bias before using a binomial test to compute enrichment *p* values. Peaks within CCANs were used as target sequences, whereas all other cluster-specific peaks were used as background regions. For downstream analysis, we took motifs that were enriched with FDR ≤ 0.05 and fold enrichment ≥ 1.2.

### *In silico* identification of regulatory variants

The R package atSNP was used to quantify potential regulatory variants *in silico*^49^. atSNP takes as input a list of position probability matrices and a list of SNP loci and uses importance sampling to assess the significance of an observed TFBS affinity change when the SNP allele is introduced. Our input set of TFBSs and SNPs passed to atSNP was based on the following criteria:

Open chromatin region criteria:

a) TFBS-SNP pair was located within a DAP
b) TFBS-SNP pair was located within a unidirectional or mixed CCAN
c) CCAN contained ≥ 1 peak that was within 500 Kbp from a GWAS SNP
d) DAP containing the TFBS was highly coaccessible (coaccessibility score of ≥ 0.2) with a peak overlapping the promoter or intron 1 of a DEG in the closest matching snRNA cluster
e) DAP and associated DEG had the same sign log fold change

TFBS criteria:

f) Motif *p* value at the TFBS was ≤ 5E-5 (for major *or* minor allele sequence)
g) TFBS overlapped ≥ 1 SNP SNP

criteria:

h) Overlapping SNP had a minor allele frequency ≥ 1%. For multiallelic SNPs, only alleles with frequency ≥ 1% were included.

atSNP assumes the underlying nucleotide distribution follows a first order Markov model. The sequences in the proposal distribution are of length 2*L*-1, and contain a subsequence of length *L*, which matches the motif in question. The sequences are weighted based on the affinity score of the matching subsequence and the expected affinity change resulting from a single nucleotide alteration at the SNP position, *L*. atSNP outputs a “1” in the event that a selected sequence has an expected affinity change greater than or equal to that which was observed, and a “0” otherwise. This number is multiplied by the sample weights, i.e. the null and proposal distributions’ likelihood ratio. The mean value after *N* Monte Carlo samples is taken as the estimated *p* value, where *N* is determined by the length of the motif. In addition, we ran 1000 iterations of atSNP for every SNP-TFBS overlap to further reduce the variance estimate from the Monte-Carlo sampling procedure. We reported the mean *p* value, variance, minimum and maximum *p* value for each atSNP test across all 1000 iterations. In reporting key results, we controlled the FDR at 0.05 using the Benjamini-Hochberg procedure.

Our flagship results included in the main/supplemental figures passed the following additional criteria:

a. Dysregulated TF showed expression in ≥10% of cells in the corresponding snRNA cluster
b. Target DEG associated with the regulatory SNP had log fold change magnitude of ≥ 0.15
c. FDR ≤ 0.01

## DECLARATIONS

### Ethics approval and consent to participate

The project was approved by the Duke Institutional Review Board (IRB). The study does not involve living human subjects. All samples were obtained from autopsies, and all are de-identified.

### Consent for publication

Not applicable.

### Availability of data and materials

The single-nucleus (sn)RNA-Sequencing and snATAC-Sequencing data will be available at the Synapse data repository (https://synapse.org, ProjectSynID: syn50996869). Access will be avaliable upon publication under controlled use conditions. In addition, the snRNA-seq and snATAC-seq raw and normalized count data generated in this study will be available at the Duke Research Data Repository (https://research.repository.duke.edu). All computer code used for this study will be available before publication on GitHub (https://www.github.com).

### Authors’ contributions

O.C-F. conceived of the presented idea. J.G. and O.C-F. acquired brain samples and designed the study sample. J.C., D.G. and J.B. performed tissue processing, nuclei extraction, and the 10x experiments for snATAC-seq/snRNA-seq library preps. J.G., D.G. and E.K.S performed all data analyses. M.E.G. performed covariate analysis. C.H. performed DEG annotation. O.C-F. planned and supervised the work. G.E.C. and A.A.K. provided guidance. J.G., D.G., E.K.S., M.E.G., G.E.C., A.A.K., and O.C-F. discussed and interpreted the results. J.G., D.G. and E.K.S. prepared figures and tables. J.G., E.K.S. and O.C.-F. wrote the manuscript. D.G., M.E.G., G.E.C., and A.A.K. reviewed and edited the manuscript. All authors read and approved the final manuscript. O.C-F. obtained funding support.

## Acknowledgements

We thank the Kathleen Price Bryan Brain Bank at Duke University (funded by NIA <u>AG028377</u>) for providing us with the brain tissues, and the Duke Sequencing and Genomic Technologies Shared Resource for sequencing. We thank Dr. M. Lutz for generously providing his advice and guidance on bioinformatics resources. This work used a high-performance computing facility partially supported by grant 2016-IDG-1013 (“HARDAC+: Reproducible HPC for Next-generation Genomics”) from the North Carolina Biotechnology Center.

---

**Figure S1.**
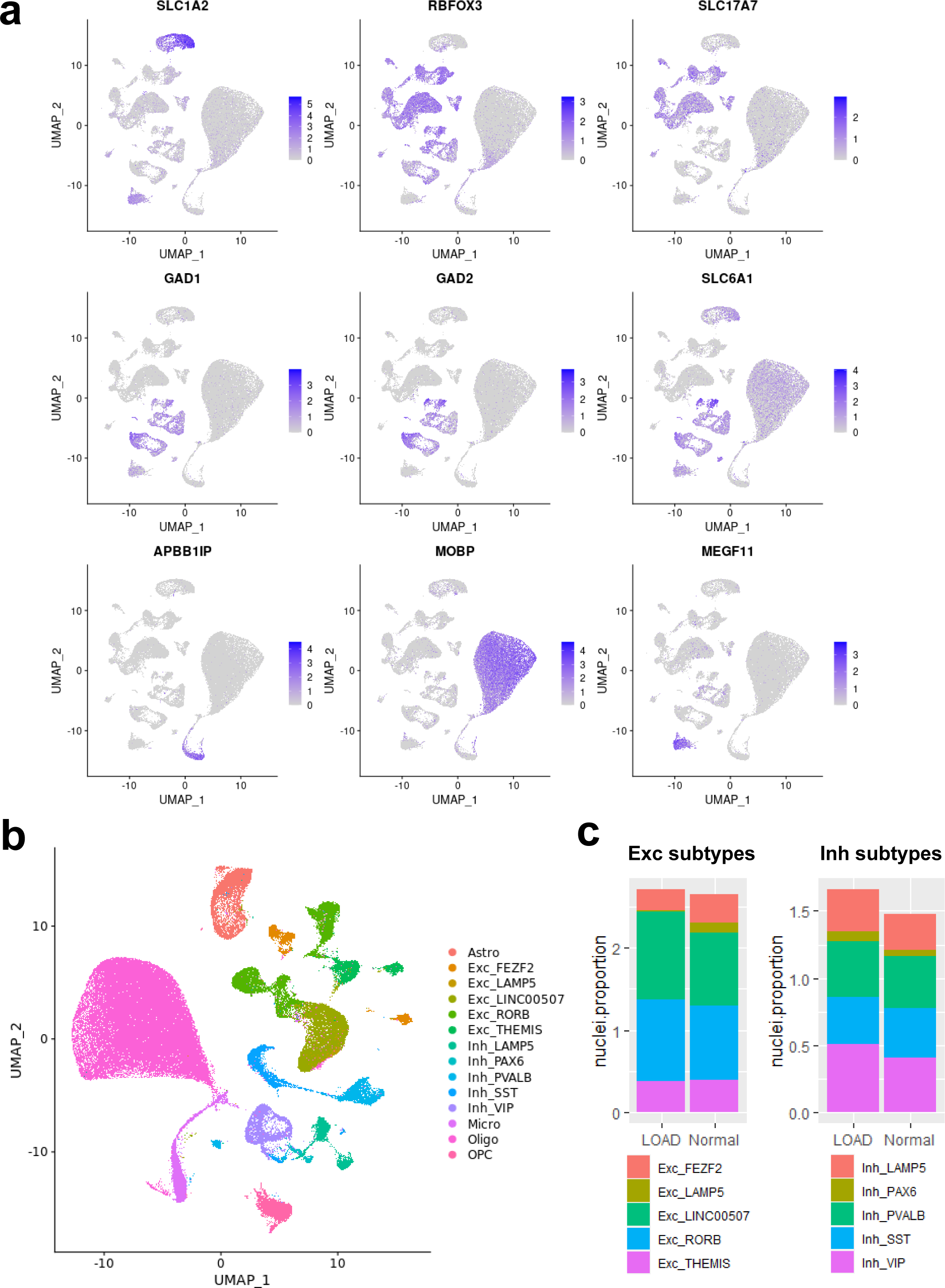
Confirmation of cluster annotation. **a,** Feature plots of log-normalized, corrected count data from SCTransform output showing cell type-specific markers for astrocytes (*SLC1A2*), neurons (*RBFOX3*), excitatory neurons (*SLC17A7*), inhibitory neurons (*GAD1, GAD2, SLC6A1*), microglia (*APBB1IP*), oligodendrocytes (*MOBP*), and OPCs (*MEGF11*). **b**, UMAP plot of re-annotated snRNA-seq dataset with known excitatory and inhibitory neuron subtypes. **c**, Proportions of nuclei of each of the five subtypes of excitatory and inhibitory neurons for each sample. Average proportions for the 12 LOAD and 12 Normal samples are shown.

**Figure S2.**
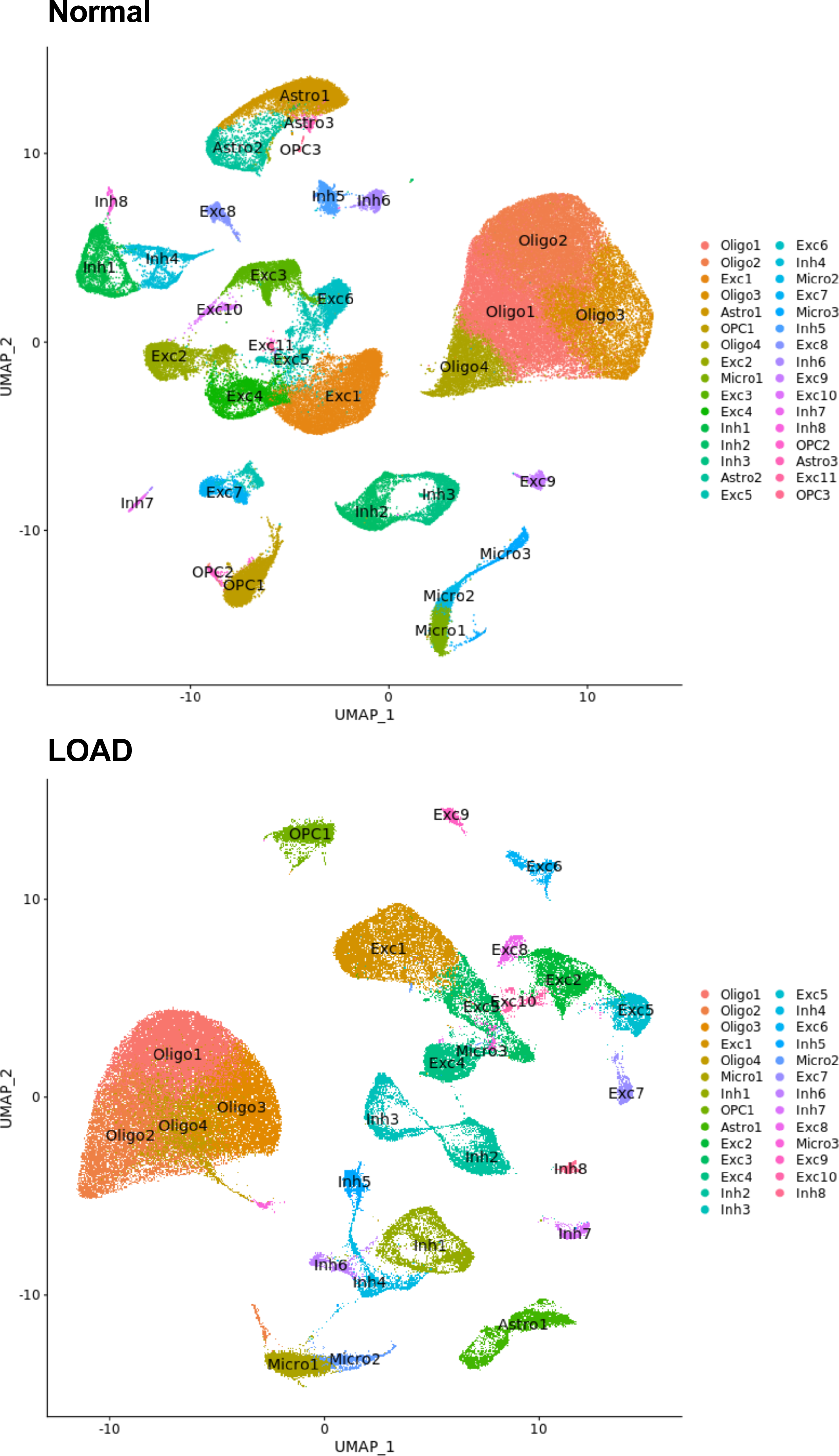
Cell subtype UMAP clustering of separately integrated normal and LOAD sample sets.

**Figure S3.**
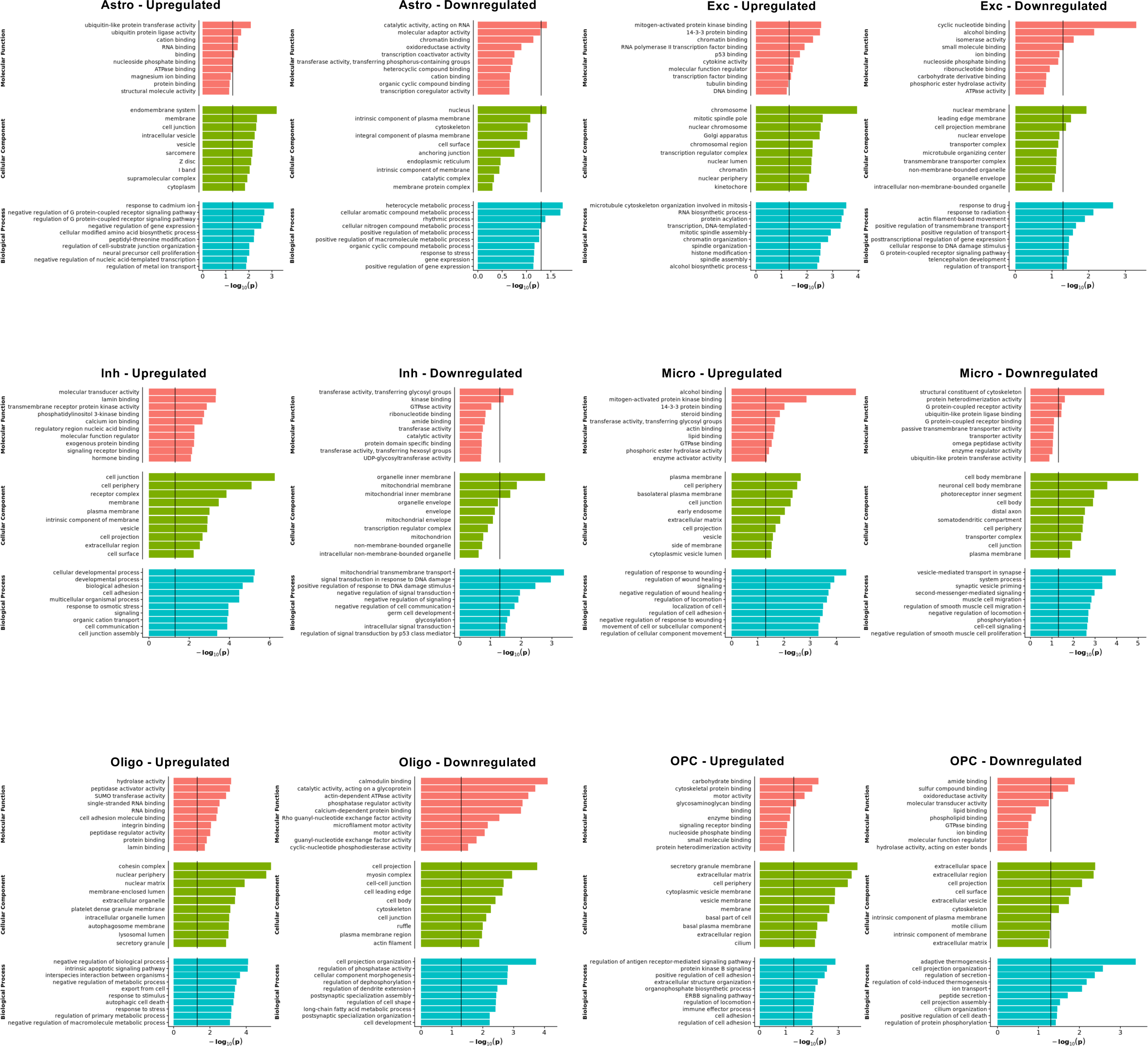
Gene ontology analysis of DEGs identified in snRNA-seq data for cell types.

**Figure S4.**
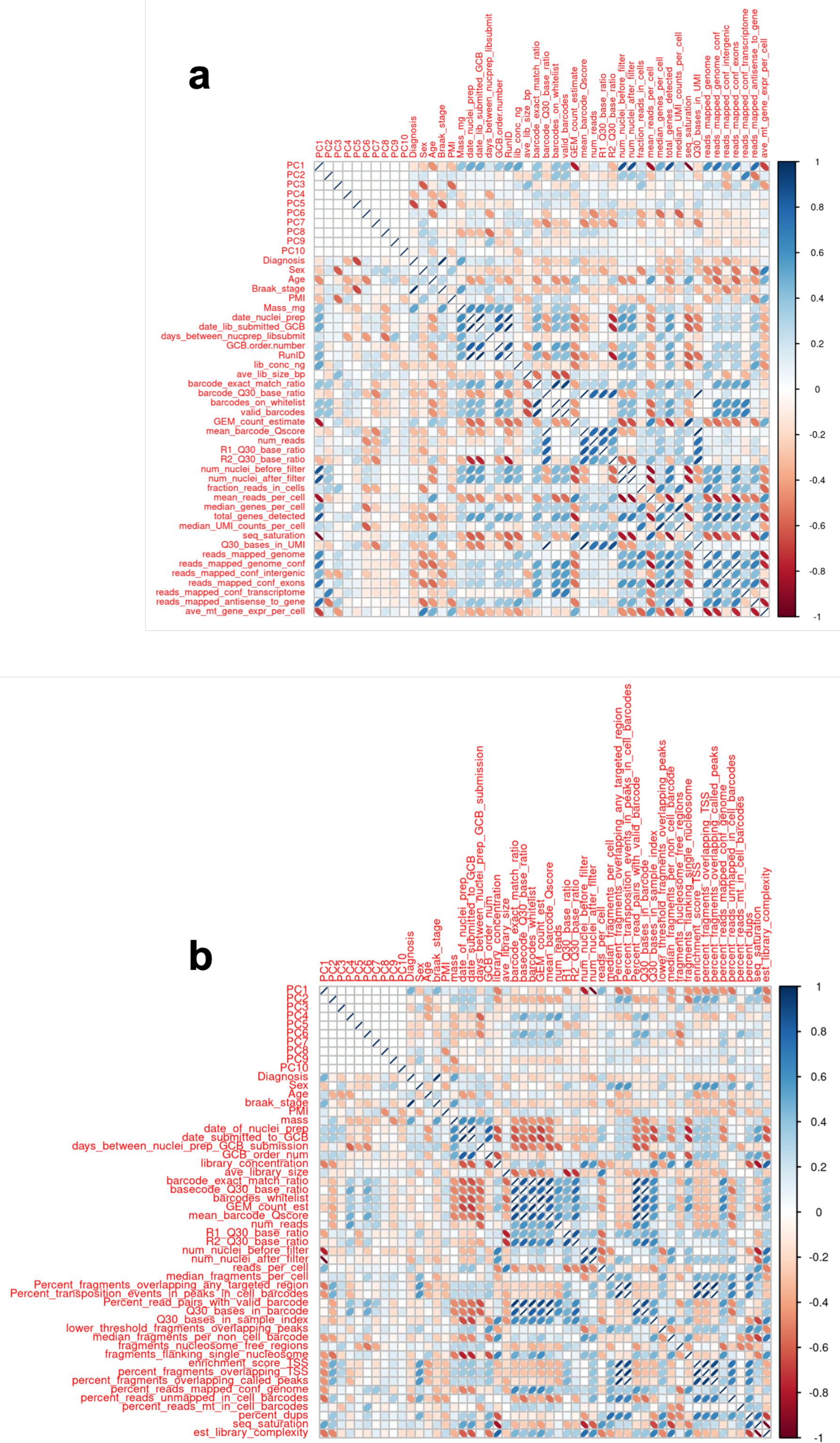
Correlation of metadata covariates in snRNA-seq and snATAC-seq data.

**Figure S5.**
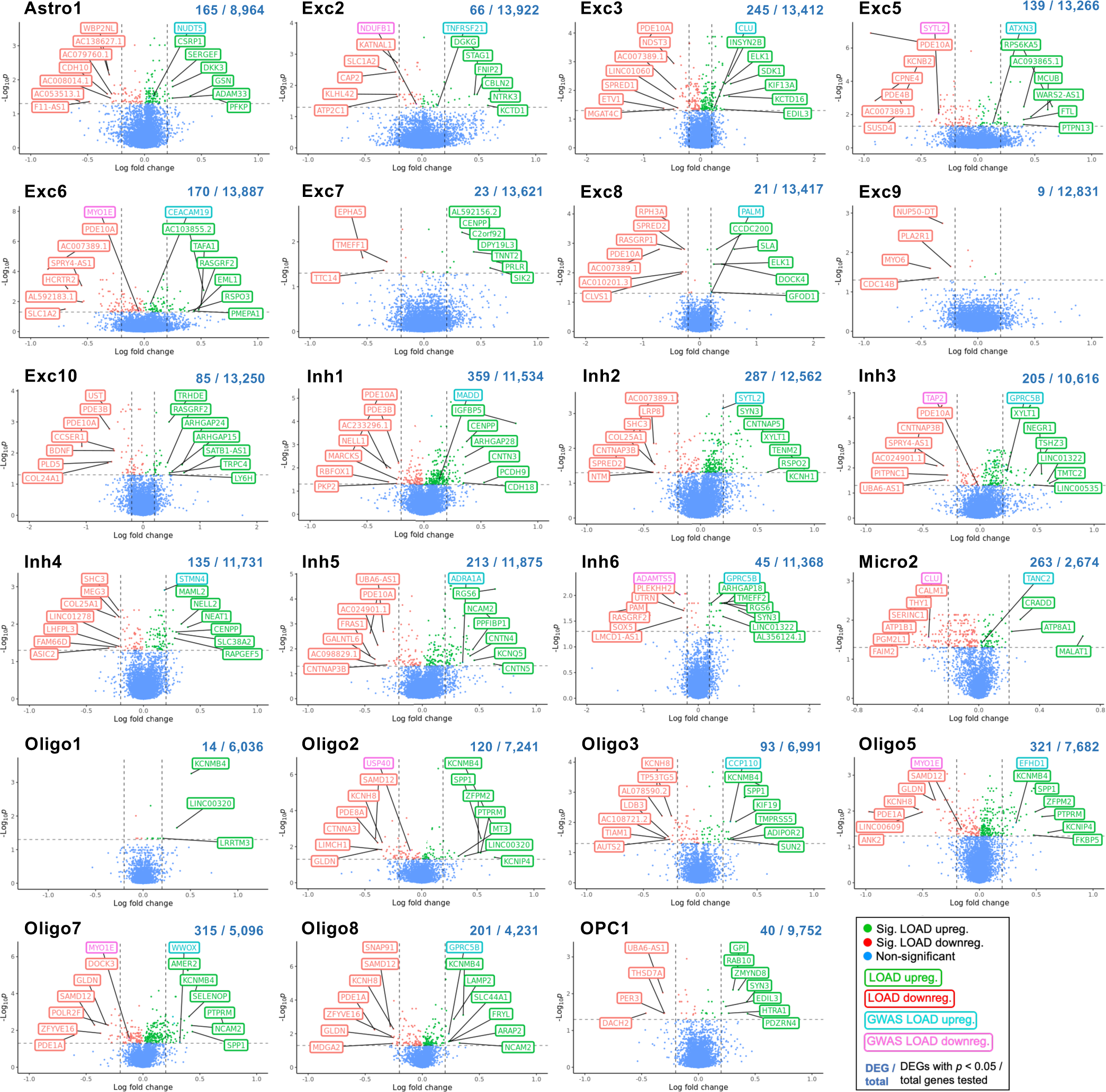
Top differentially-expressed genes (DEGs) upregulated and down-regulated in LOAD by cluster. Unbiased volcano plots for all clusters containing DEGs not shown in figure 3, representing astrocyte (Astro), excitatory neuron (Exc), inhibitory neuron (Inh), microglia (Micro), oligodendrocyte (Oligo), and oligodendrocyte precursor (OPC) cell types. Log_2_ fold change (FC) between LOAD and normal control samples is plotted against –log_10_ *p*-value (FDR). Points representing DEGs with statistically significant (*p* < 0.05) upregulation in LOAD are shown in green while DEGs with significant downregulation are shown in red. Genes without significantly differential expression are shown in blue. The proportion of DEGs to total genes examined is shown above each plot. The six DEGs with the highest absolute fold change (log_2_ FC > 0.2) in the up- and downregulated categories are labeled in green and red, respectively. The top up- and downregulated DEGs within 500kb of disease-associated SNPs previously identified in GWAS are labeled in teal and pink, respectively. The Inh8 and Inh9 clusters did not contain DEGs and are not shown.

**Figure S6.**
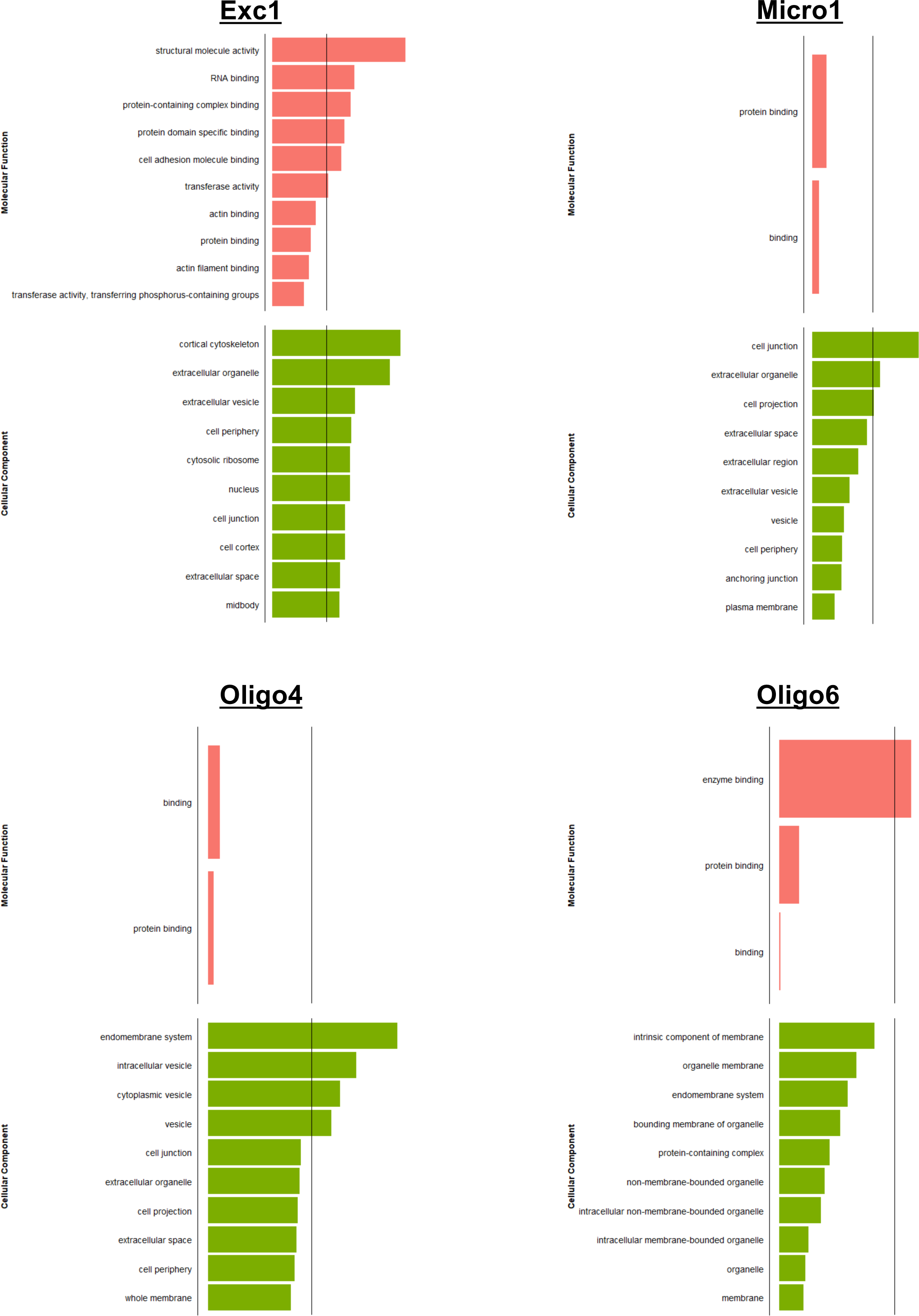
Additional gene ontology of cCRE-linked DEGs. Gene ontological analysis of molecular functions and cellular components for cCRE-linked DEGs associated with CCANs in the indicated cell subtype clusters. Up to the top ten enriched terms involving a minimum of three DEGs are listed.

**Figure S7.**
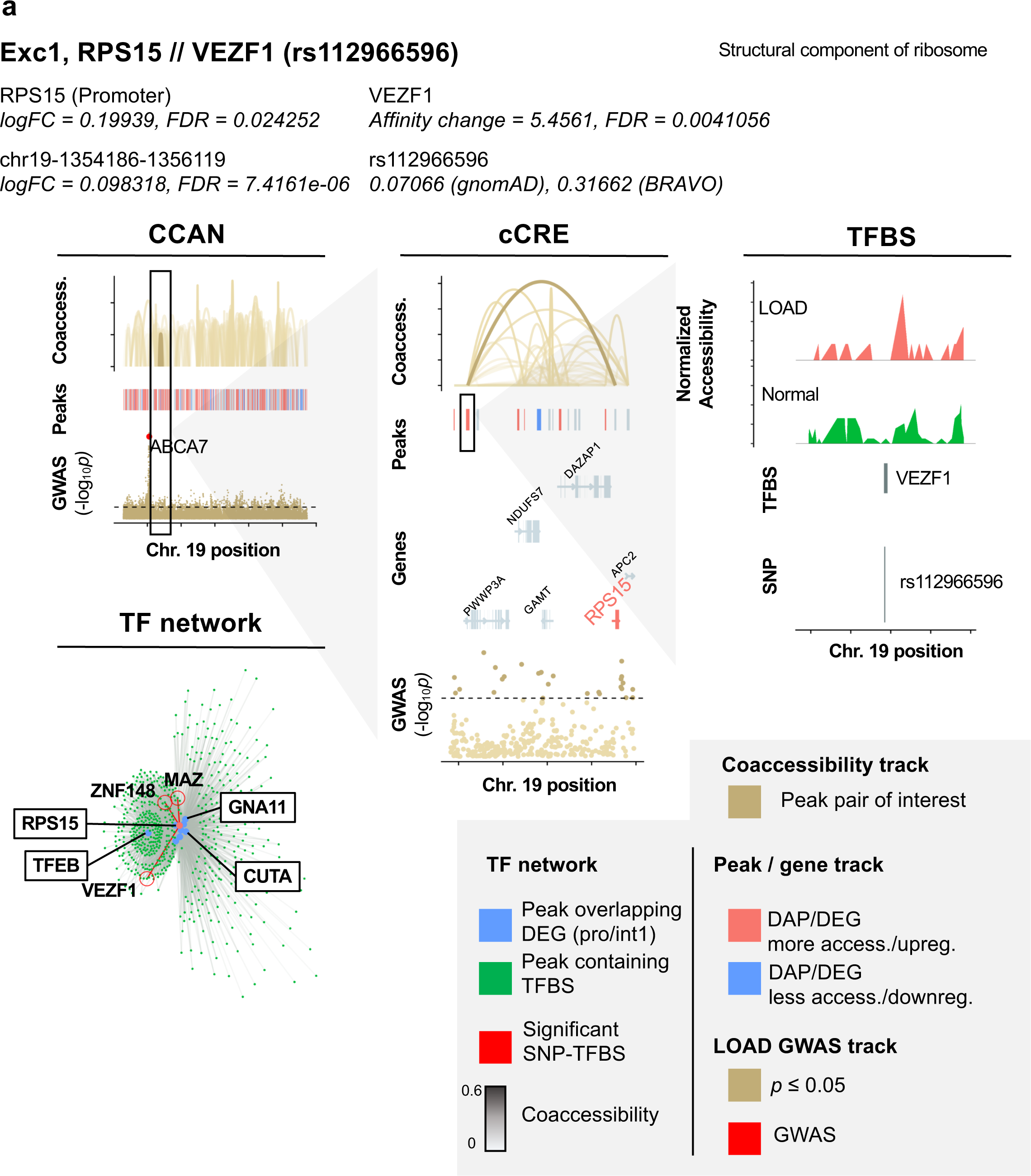
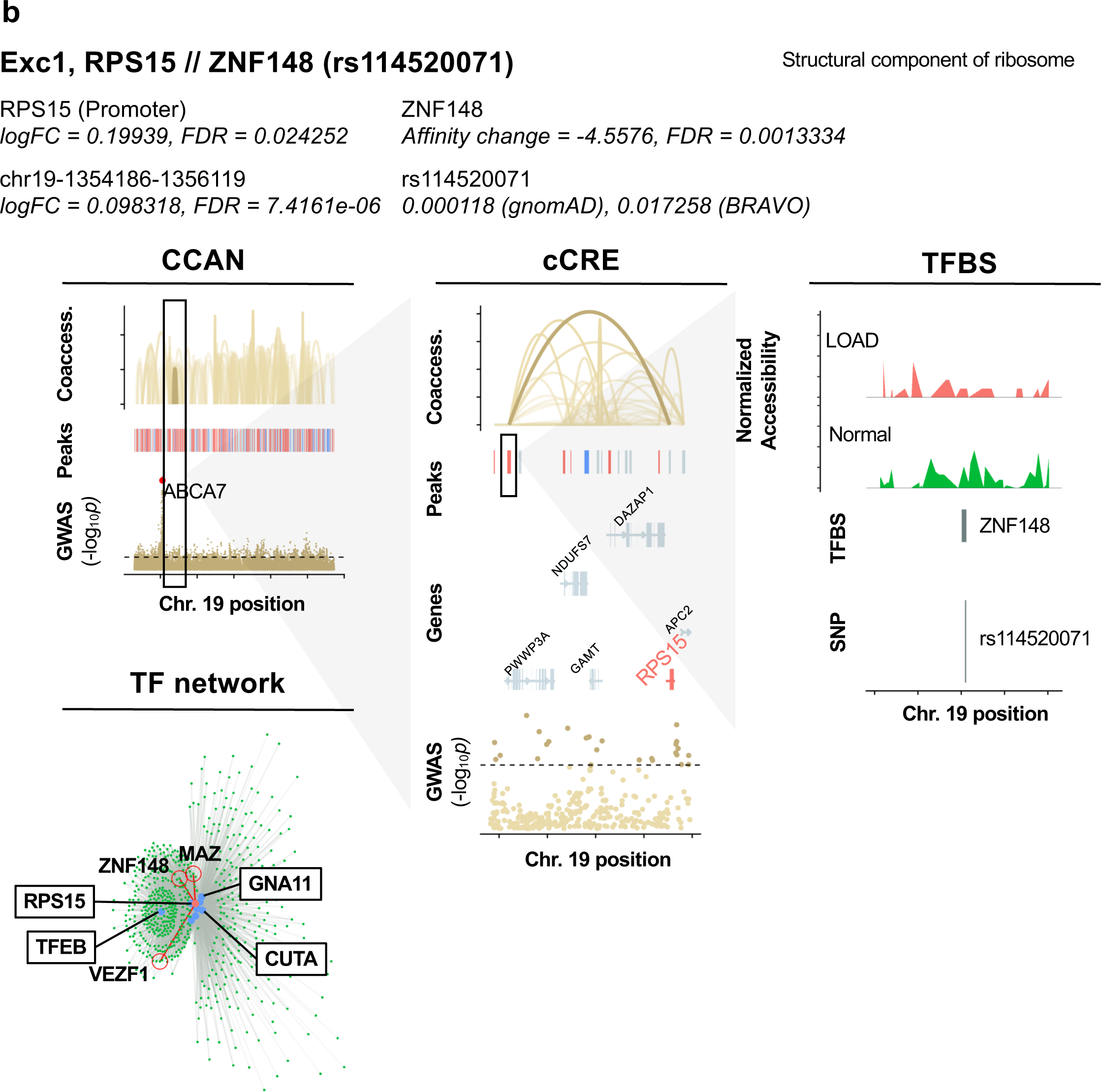
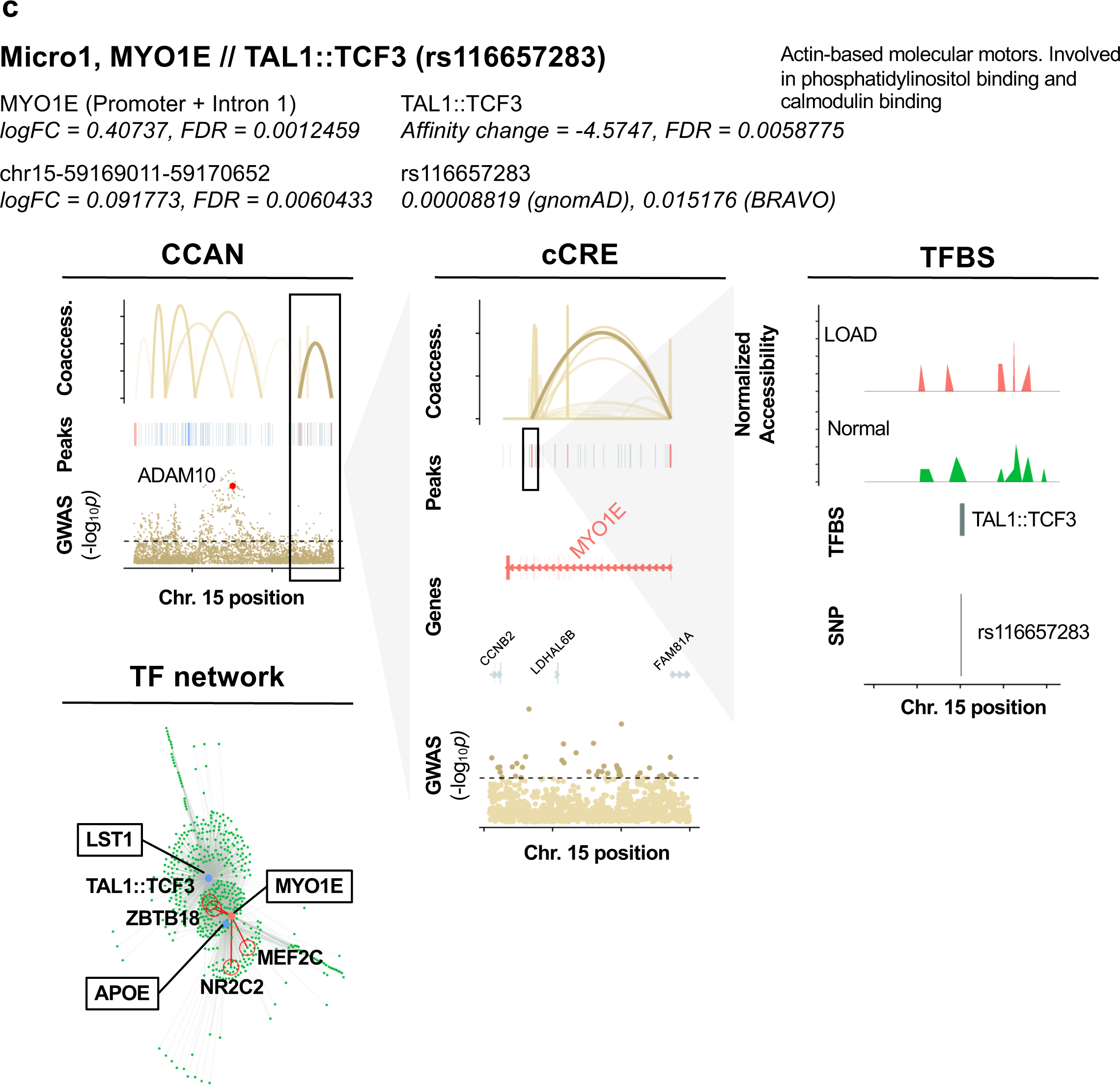
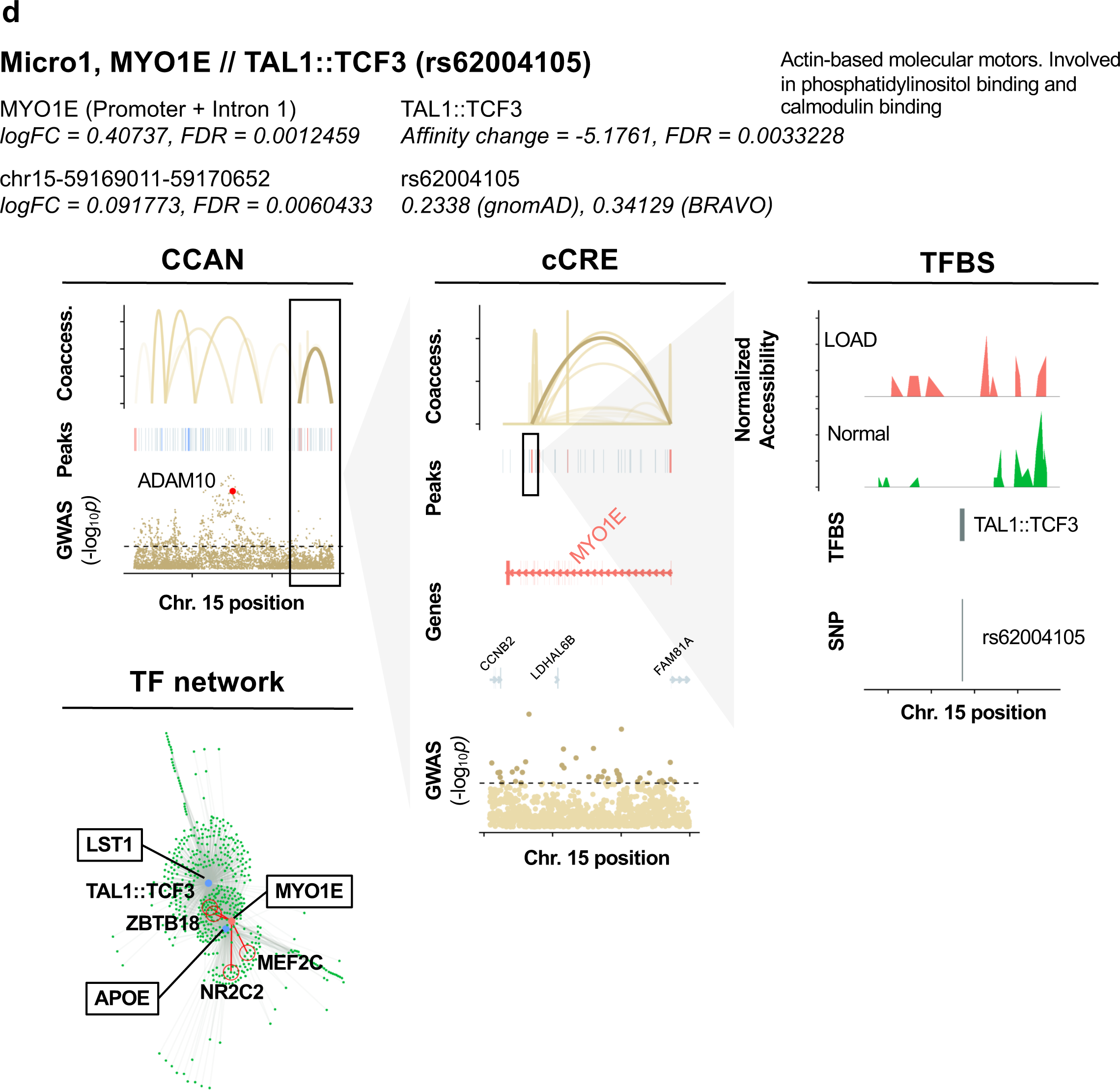
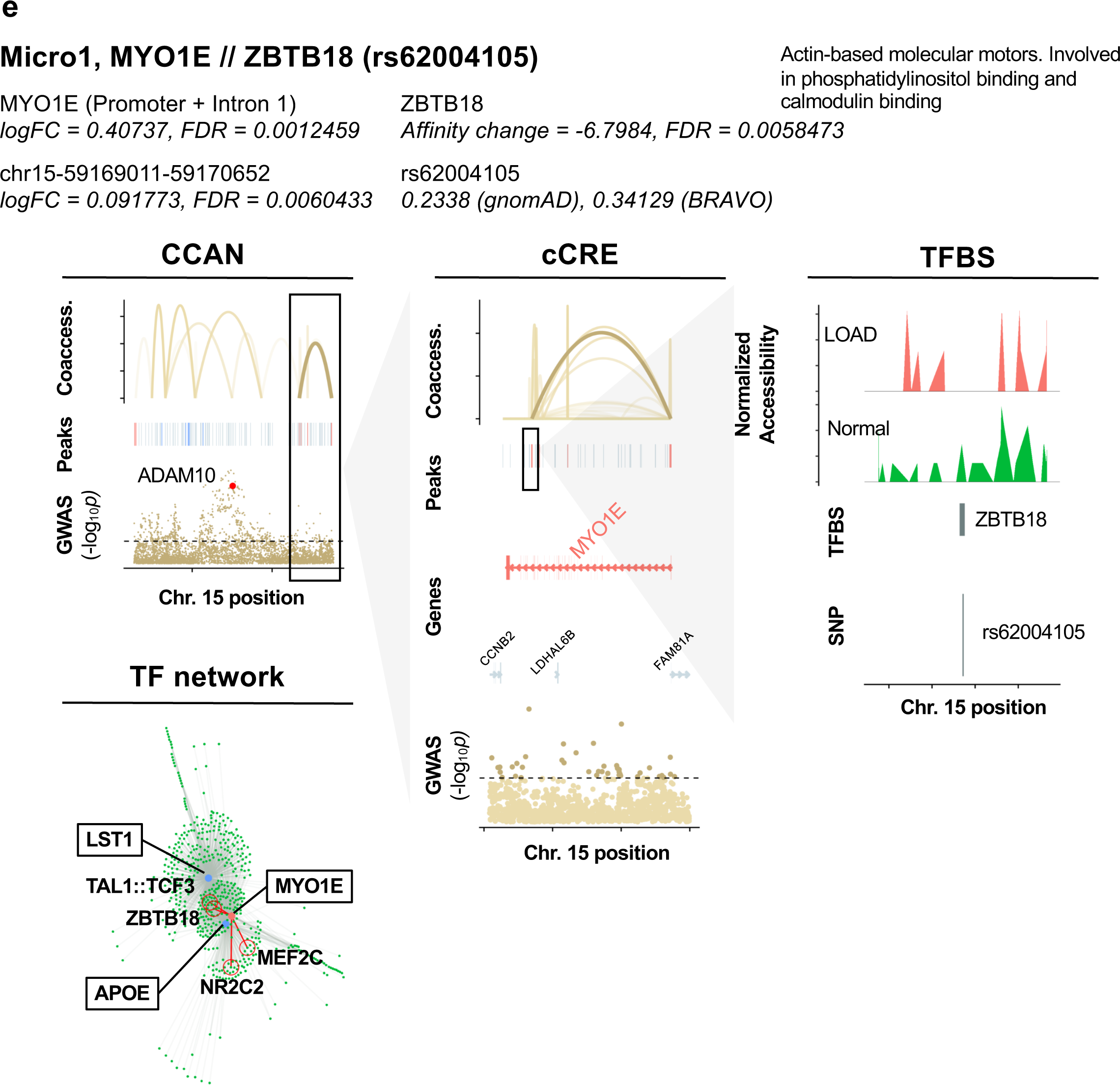
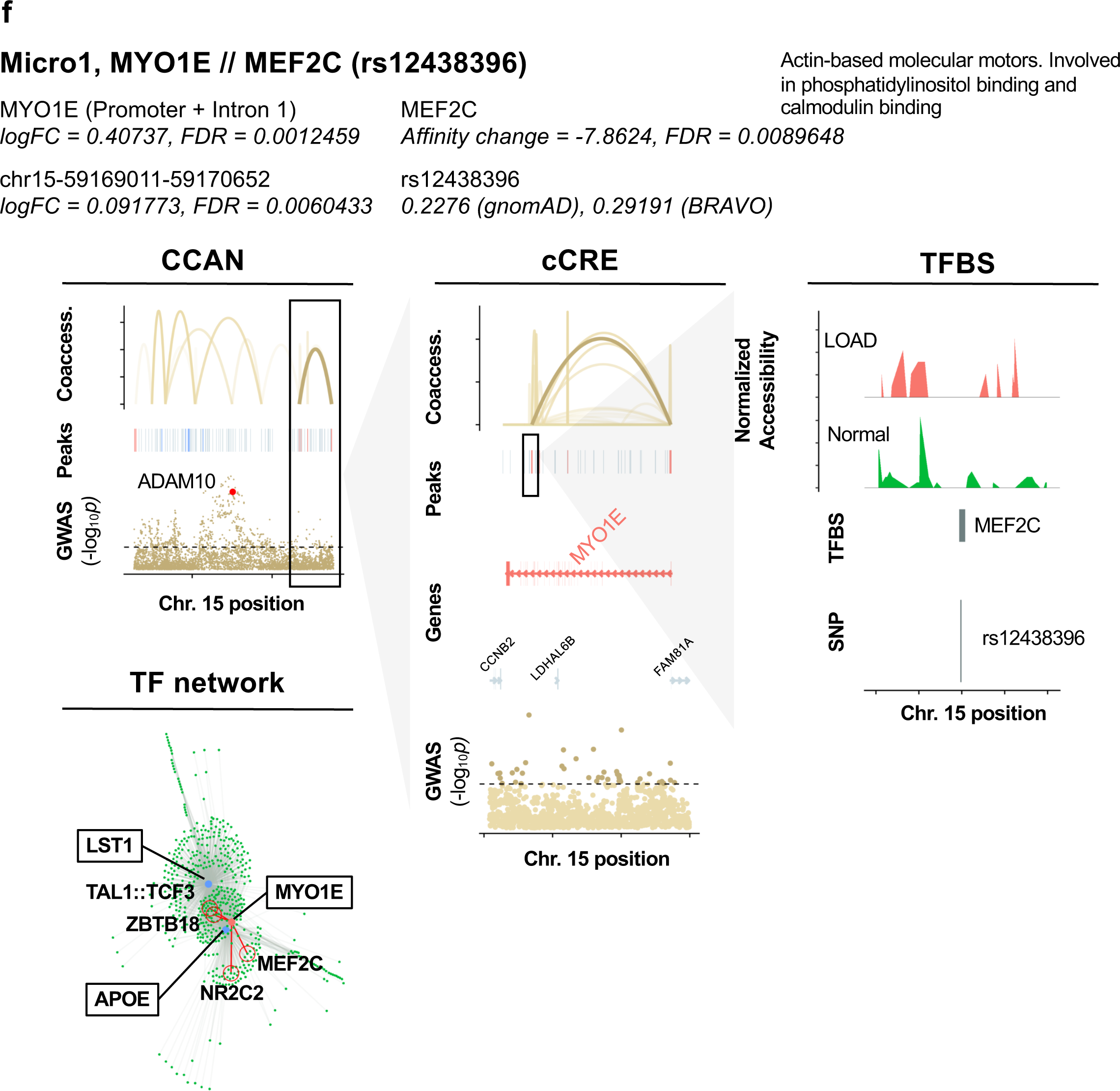
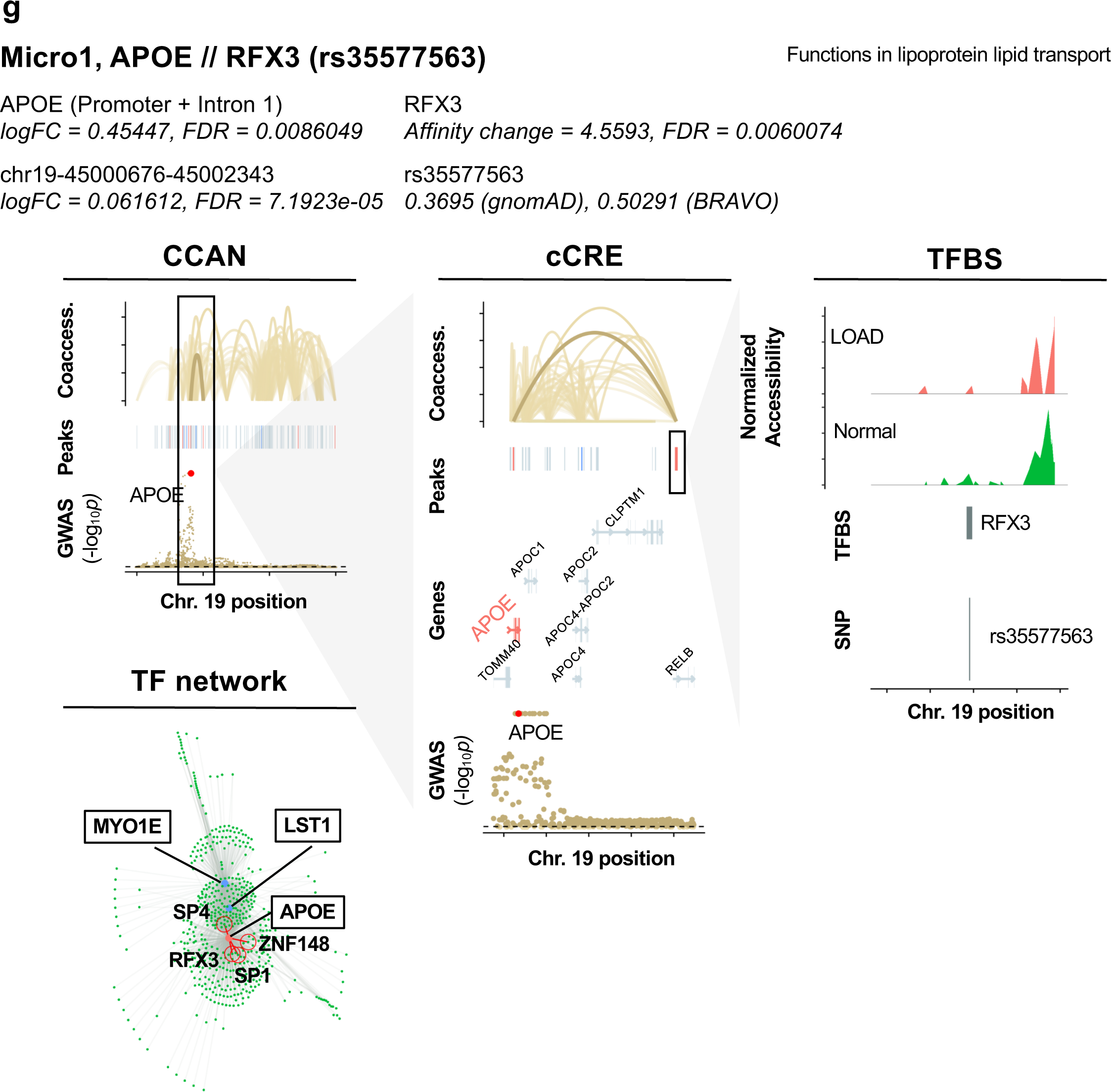
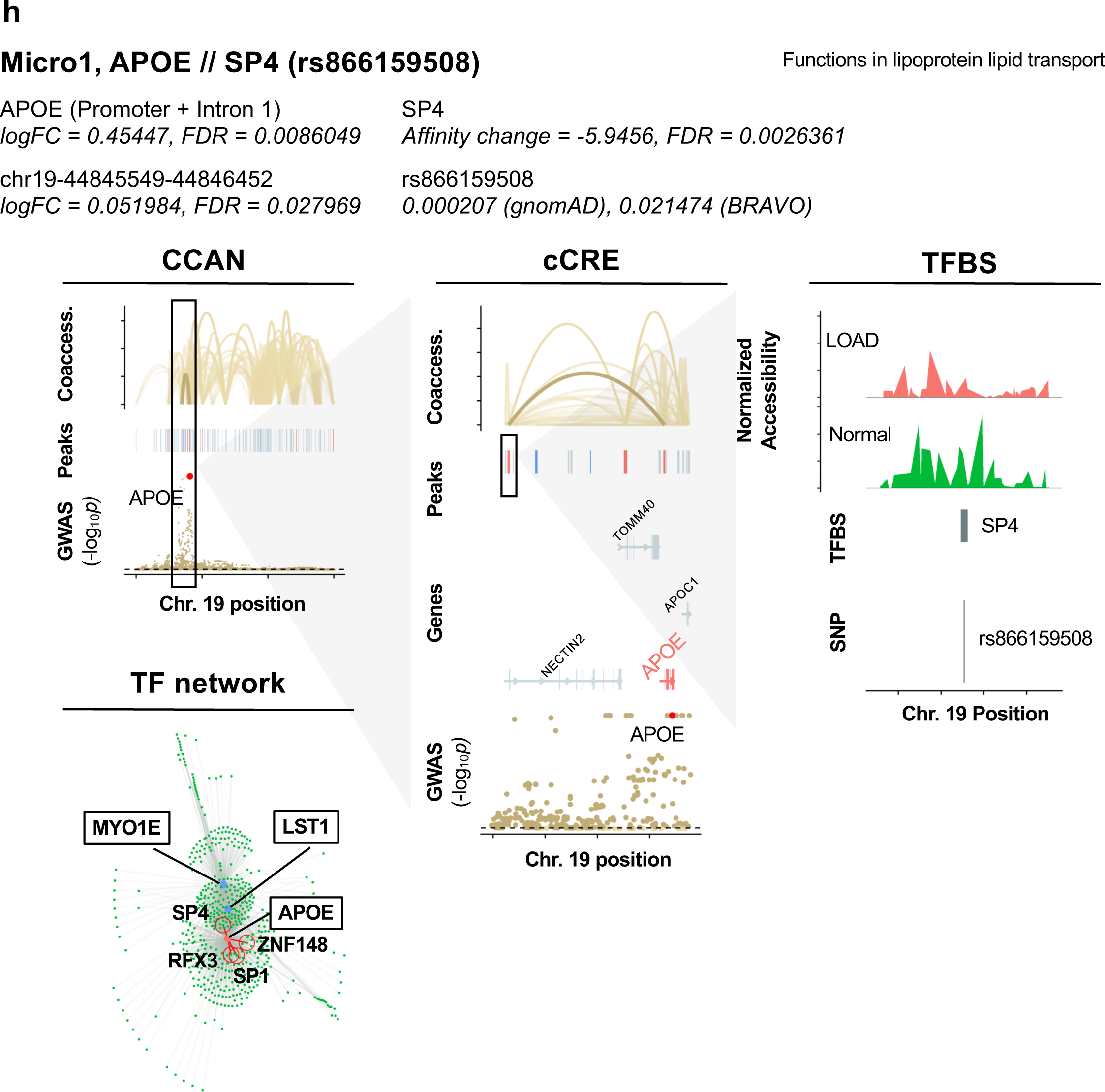
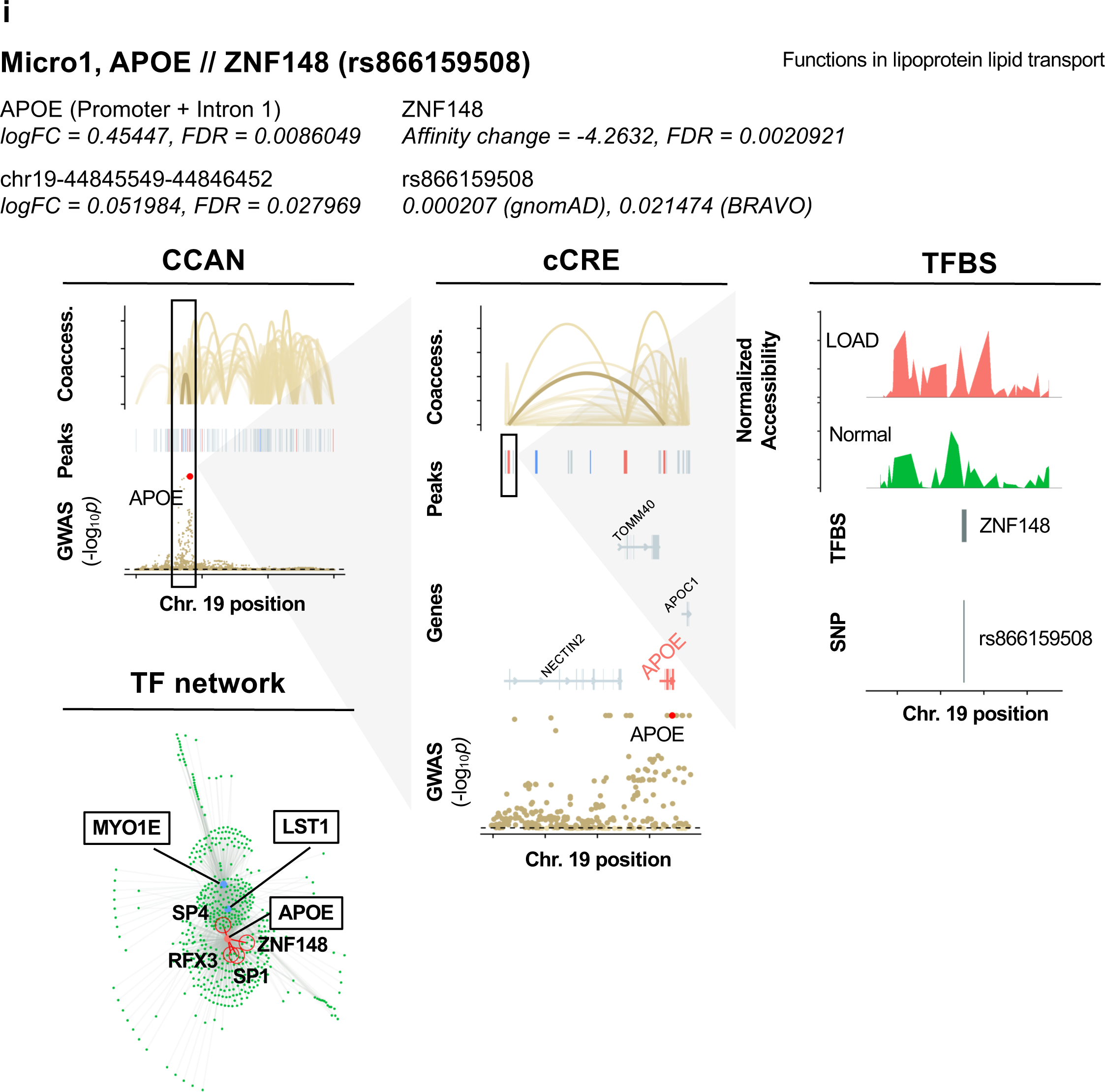
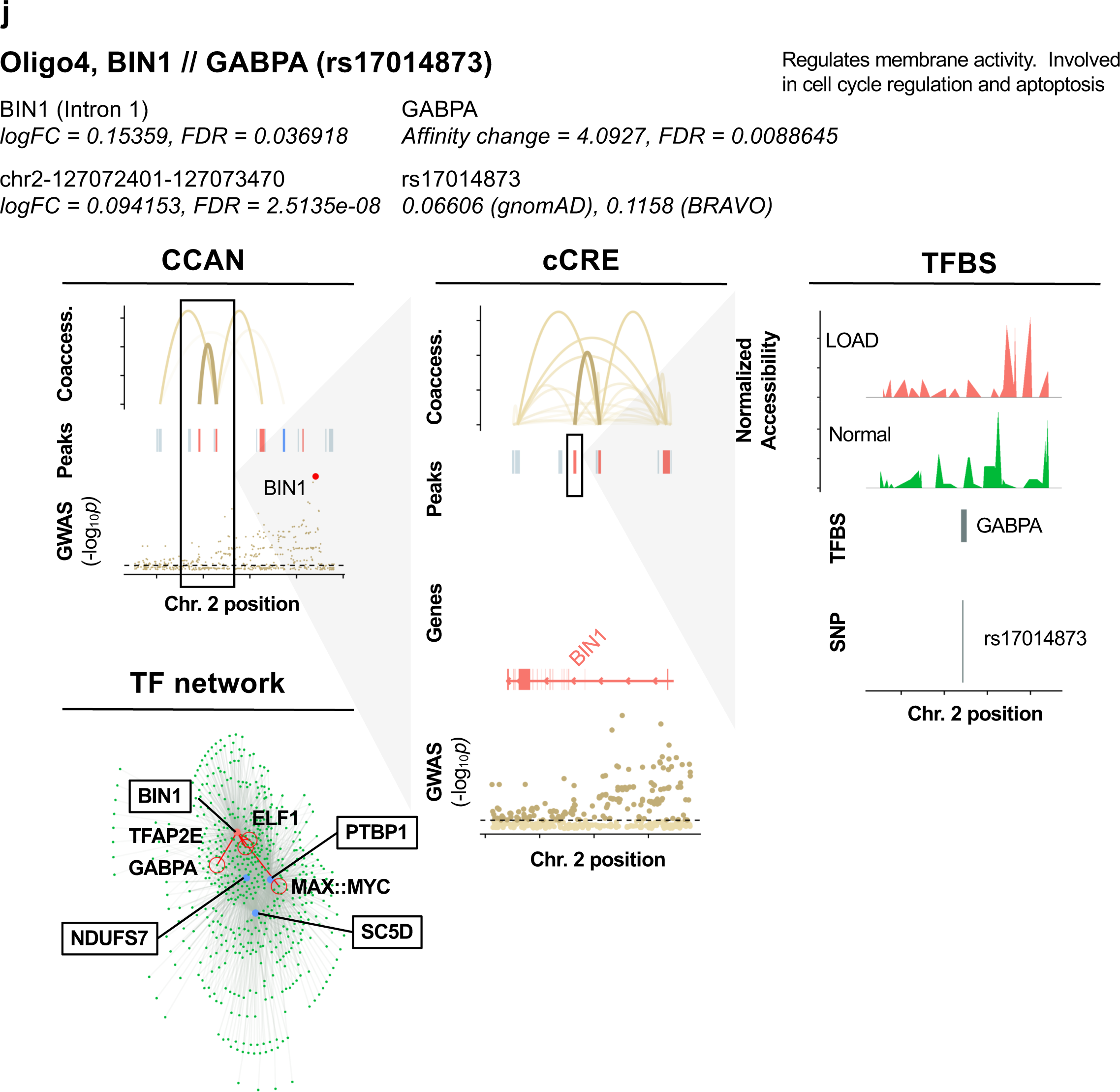
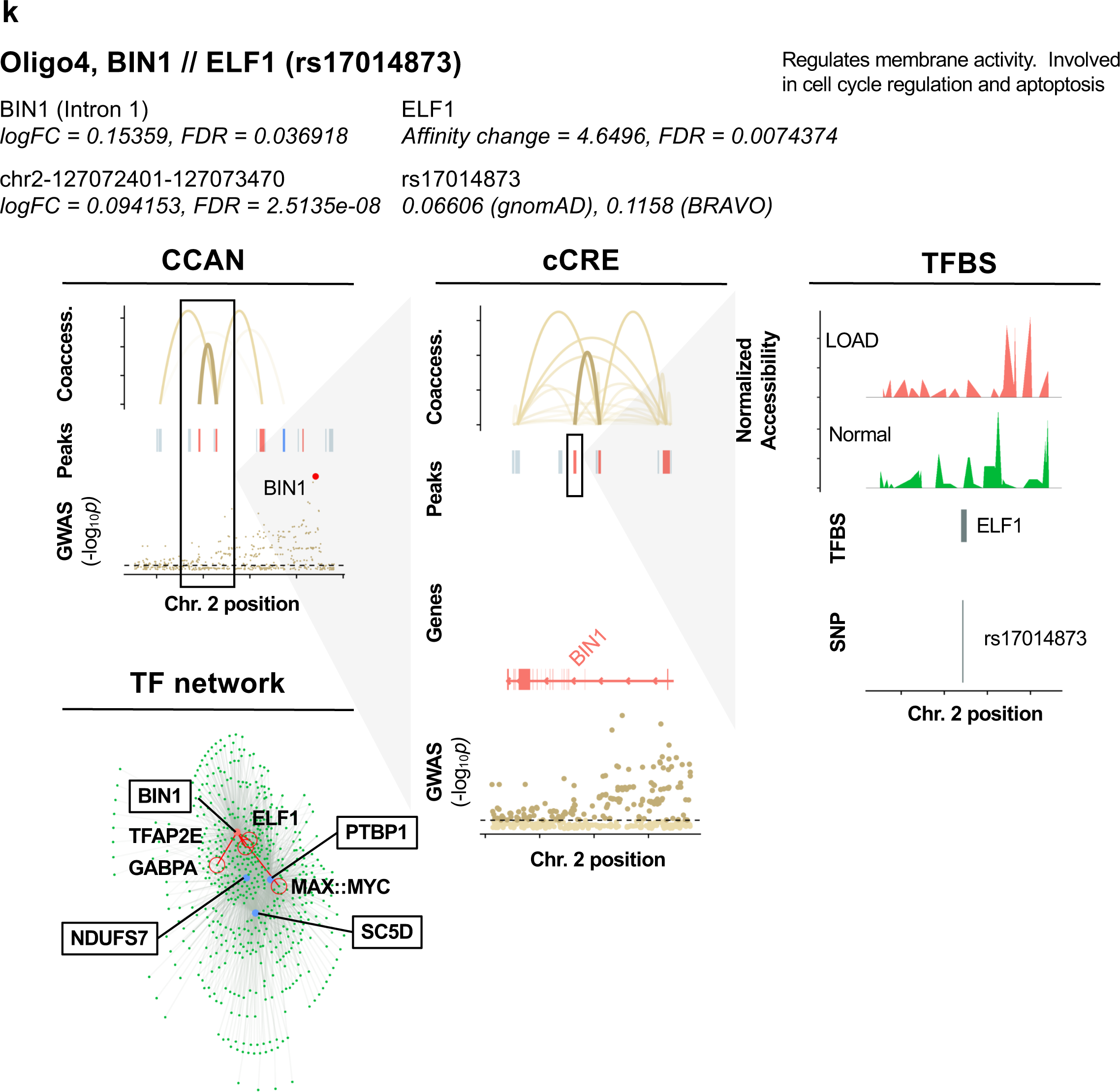
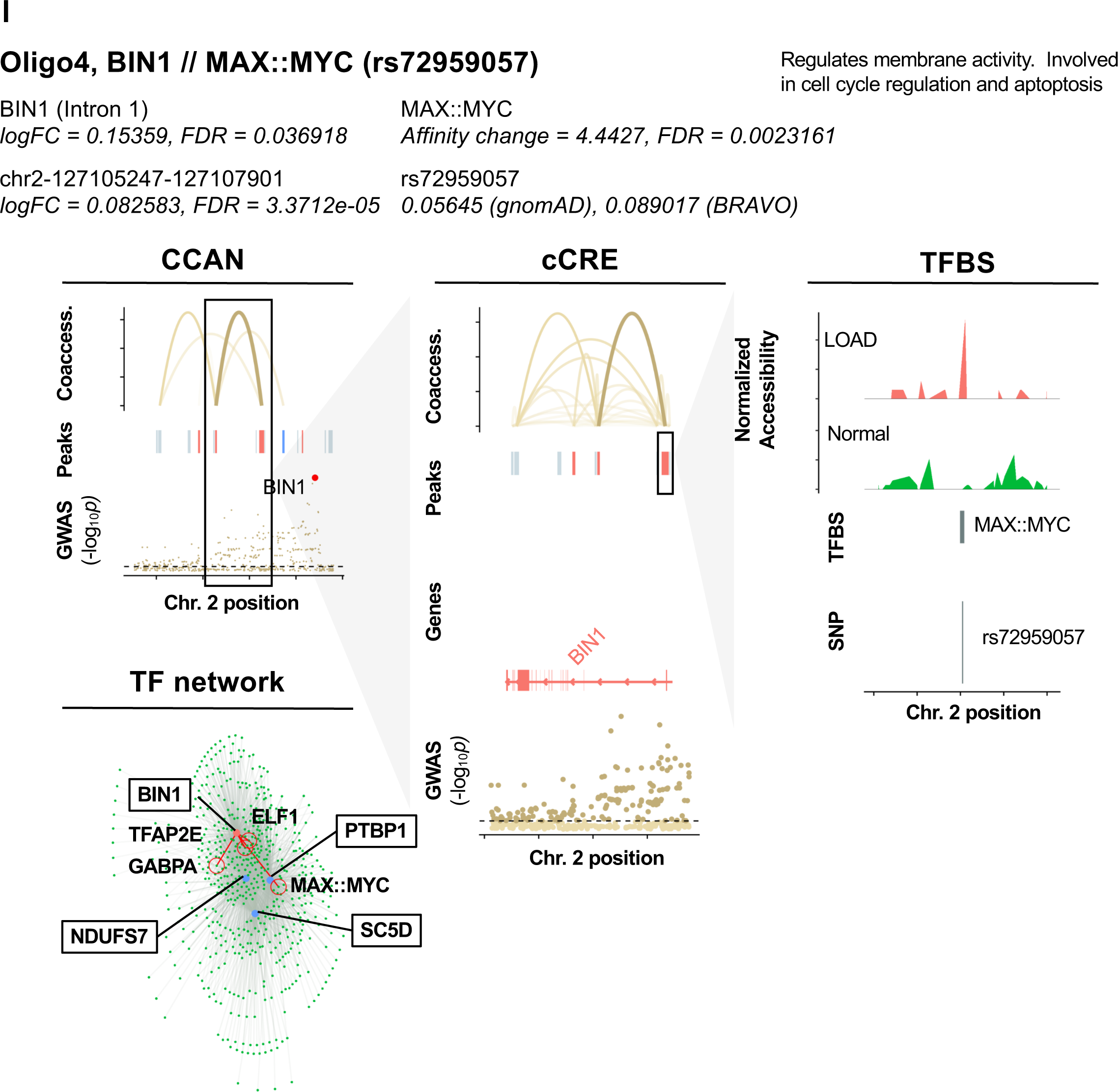
Identification of SNPs predicted to influence TF binding affinity at GWAS loci in LOAD CCANs. Diagrams of specific example SNP-TFBS overlaps. The cell subtype, regulated DEG, TF and SNP ID are shown in bold. The log fold change (*Log2FC*) and probability values (*p*) are shown for each DEG and corresponding cCRE. Additionally, functional information for each DEG is provided. CCAN stacked plots show peak coaccessibility scores, directionality of changes in DAP accessibility in LOAD (red = increased accessibility, blue = reduced accessibility), and degree of LOAD association for GWAS loci. All features are arranged along the same horizontal access to indicate chromosomal position. cCRE stacked plots are detailed from boxed area of CCAN plots additionally indicate overlapped gene coding regions, with upregulated DEGs shown in red and downregulated DEGs shown in blue. TFBS activity stacked plots are detailed from boxed areas of cCRE plots and indicate normalized accessibility of genomic region in LOAD and normal samples as well as aligned chromosomal positions of TFBSs and SNPs. TF Network plots illustrate potential regulatory networks between DEG-overlapping peaks (blue) and TFBS-overlapping peaks (green), with those linkages predicted to be affected by LOAD SNPs shown in red.

